# Pan-Cancer analysis of somatic mutations in miRNA genes

**DOI:** 10.1101/2020.06.05.136036

**Authors:** Martyna Olga Urbanek-Trzeciak, Paulina Galka-Marciniak, Paulina Maria Nawrocka, Ewelina Kowal, Sylwia Szwec, Maciej Giefing, Piotr Kozlowski

**Affiliations:** Institute of Bioorganic Chemistry, Polish Academy of Sciences, Poznan, Poland; Institute of Human Genetics, Polish Academy of Sciences, Poznan, Poland

**Keywords:** miRNA, somatic mutations, Pan-Cancer, TCGA, noncoding

## Abstract

miRNAs are considered important players in oncogenesis, serving either as oncomiRs or suppressormiRs. Although the accumulation of somatic alterations is an intrinsic aspect of cancer development and many important cancer-driving mutations have been identified in protein-coding genes, the area of functional somatic mutations in miRNA genes is heavily understudied. Here, based on analysis of the whole-exome sequencing of over 10,000 cancer/normal sample pairs deposited within the TCGA repository, we identified and characterized over 10,000 somatic mutations in miRNA genes and showed that some of the genes are overmutated in Pan-Cancer and/or specific cancers. Nonrandom occurrence of the identified mutations was confirmed by a strong association of overmutated miRNA genes with KEGG pathways, most of which were related to specific cancer types or cancer-related processes. Additionally, we showed that mutations in some of the overmutated genes correlate with miRNA expression, cancer staging, and patient survival. Our results may also be the first step (form the basis and provide the resources) in the development of computational and/or statistical approaches/tools dedicated to the identification of cancer-driver miRNA genes.

## Introduction

Cancer encompasses a broad spectrum of heterogeneous diseases whose development (i.e., initiation, promotion, and progression) is associated with the accumulation of numerous genetic alterations in the cancer genome, which is the hallmark of all cancers. Although most of these alterations are neutral, some of the randomly occurring mutations are functional, providing a growth advantage to a neoplastic cell ^1^. As a result of clonal selection of the fastest dividing cells, functional (driver) mutations often recur in genes playing an important role in cancer development (driver genes) and therefore may serve as indicators of such genes. Numerous large cancer genome sequencing studies (mostly whole-exome sequencing, WES) have been performed, and hundreds of cancer-driving genes and thousands of cancer-driving mutations have been detected. Some of these genes/mutations, e.g., *EGFR, BRAF*, and *JAK2*, serve as important biomarkers for cancer-targeted therapies. As the overwhelming majority of the cancer genome studies have focused on protein-coding genes, the most identified cancer-driver mutations are in protein-coding sequences, which encompass barely 2% of the genome. The spectacular exception are TERT promoter mutations, which occur most frequently in melanoma, brain, and bladder cancers but have also been identified in other cancers ^2–4^.

On the other hand, a growing body of evidence indicates that miRNAs, a class of short (~21 nt long) single-stranded noncoding RNAs, play an important role in cancer, and it was shown that particular miRNAs can either drive (oncomiRs, often upregulated in cancer) or suppress (suppressormiRs, often downregulated in cancer) oncogenesis. It was also proposed that miRNAs have great potential as cancer biomarkers and/or targets of cancer therapies ^5–8^.

Among the most intensively studied miRNAs whose function in cancer is best documented are the let-7 family, miR-17-92 cluster (oncomiR-1), miR-21, and miR-205 (reviewed in ^9^). Although the global level of miRNA is generally downregulated in cancer, many miRNAs are consistently either upregulated or downregulated in particular cancer types or specific cancer conditions. It was also shown that miRNA genes frequently show copy number alterations (either amplification or deletion) in cancer ^10,11^. The cancer-related processes that are regulated by miRNAs include cell proliferation, epithelial-mesenchymal transformation (EMT), migration, angiogenesis, inflammation, apoptosis, and response to cancer treatment (reviewed in ^12–15^).

Despite the great interest in the role of miRNA in cancer, very little (close to nothing) is known about somatic mutations in miRNA genes (defined here as sequences coding for the most crucial part of miRNA precursors) occurring in cancer. Considering subsequent steps of miRNA biogenesis and the mechanism of miRNA posttranscriptional gene regulation, mutations may be expected to affect different attributes of miRNA genes. In addition to the most obvious consequences of mutations in seed sequences that affect the pivotal function of miRNAs, i.e., the ability to recognize and downregulate their specific targets, mutations in any part of the miRNA precursor may affect the effectiveness or precision of miRNA biogenesis by altering/destabilizing the hairpin structure of the miRNA precursor, by altering DROSHA or DICER1 cleavage sites, or by altering protein-interacting or other regulatory sequences/structure motifs ^16–19^. Additionally, mutations destabilizing one of the miRNA duplex ends may alter 5p/3p miRNA preference. Despite the scarcity of identified miRNA gene mutations, the individual examples of SNPs, germline or somatic mutations provide proof, at least for some of the scenarios listed above. Examples include (i) the mutation in the seed sequence of miR-204-5p, affecting target recognition, that causes inherited retinal dystrophy ^20^; (ii) the mutation in the passenger strand of *hsa-miR-96* that destabilizes the structure of the miRNA precursor, affects its processing, and decreases the miR-96-5p level, eventually resulting in the same phenotypic effect as mutations in the seed sequence of the guide strand, i.e., nonsyndromic inherited hearing loss ^21,22^; (iii) the G>C substitution (SNP rs138166791) in the penultimate position of the 3p passenger strand of *hsa-miR-890* that significantly lowers the cleavage efficiency by DROSHA and consequently decreases the levels of both mature miR-890-5p and passenger miR-890-3p ^23^; (iv) the G>C substitution (SNP rs2910164) located in the 3p passenger strand of *hsa-miR-146a* that is associated with an increased risk of papillary thyroid carcinoma, where it was shown that the C allele of the SNP alters the structure of the precursor, decreases expression of the mature miRNA and activates the passenger strand, which becomes the second mature miRNA modulating many genes involved in the regulation of apoptosis ^24^; (v) interesting example is a mutation in the seed sequence of miR-184-3p, causing familial keratoconus, whose effect is not the disruption of miR-184-3p target recognition but the inability to mask overlapping targets for miR-205 in *INPPL1* and *ITGB4* ^25^; (vi) mutations in *hsa-miR-30c-1* and *hsa-miR-17* that affect the precursor structure and thereby increase the levels of mature miRNAs, downregulating *BRCA1* in familial breast cancer cases without *BRCA1/2* mutations ^26^; and finally, (vii) two different somatic mutations in the seed sequence of miR-142-3p, found in acute myeloid leukemia (AML) samples, that were shown to decrease both miR-142-5p and miR-142-3p levels and reverse the miR-5p/3p ratio (in favor of miR-3p) ^27^ (more details and references on mutations in hsa-miR-142 in the subsequent sections). An additional indication of miRNA gene sensitivity to genetic alteration is their general high conservation and the decreased density of common SNPs in miRNA hairpin sequences ^28–30^, which resembles the commonly known phenomenon of the decreased level of nonsynonymous SNPs in protein-coding sequences. In our recent analysis of somatic mutations in lung cancers, we confirmed that seed mutations affect the vast majority of the predicted targets and showed that mutations in miRNA genes often alter the predicted structure of miRNA precursors ^31^. miRNAs in cancers were also considered in the context of somatic mutations found in mRNAs that affect mRNA-miRNA and competing endogenous RNA (ceRNA)-miRNA interactions ^32,33^. As we consider sequence variations in mRNAs that may affect miRNA function, the same should apply to the sequence variations in the miRNA genes themselves.

Several multicenter projects have led to gathering data on somatic mutations from hundreds of cancers. One of the projects, The Cancer Genome Atlas (TCGA), covers over 10,000 samples from 33 types of cancers, including the most common human cancers. Importantly, the TCGA consortium works on standardized pipelines of data analysis ^34^, enabling comparisons across different cancer types (within the so-called Pan-Cancer set).

In the current study, we took advantage of data gathered by the TCGA consortium to analyze somatic mutations occurring in miRNA genes. As a result, we identified thousands of mutations in all subregions of miRNA genes and identified many Pan-Cancer or cancer-specific overmutated miRNA genes. We showed that mutations in some of the overmutated genes correlate with miRNA expression, cancer staging, and patient survival. Although the functionality of individual mutations or groups of mutations needs to be verified in independent functional studies, the strong association of the overmutated miRNA genes with cancer-related pathways indicates that miRNA gene mutations are not only random events and that at least some of them play a role in cancer.

## Methods

### Data resources

We used molecular and clinical data (Level 2) generated and deposited in the TCGA repository (http://cancergenome.nih.gov). These data included the results of somatic mutation calls in WES datasets preprocessed through the standard TCGA pipeline. We took advantage of somatic mutation data generated with four mutation caller algorithms (Mutect2, Muse, Varscan, and SomaticSnipper) and deposited as vcf.gz files. We analyzed the annotated somatic mutations with corresponding clinical information ^35^ and miRNA expression data ^36^.

### Data processing

We analyzed somatic mutations in 1918 miRNA gene regions (Supplementary Table 1) annotated in the miRBase v.22.1 database. The miRNA genes were defined as pre-miRNA-coding sequences, extended upstream and downstream by 25 nucleotides. The pre-miRNA-coding sequences were reconstructed based on 5p and 3p mature miRNA sequences defined in miRBase (in cases when only one miRNA strand was indicated, the other pre-miRNA end was reconstructed assuming the pre-miRNA hairpin structure with a 2-nt 3p overhang). According to the number of reads reported for the particular pre-miRNA arm (miRBase), the analyzed precursors were classified into one of 3 categories: (i) generating mature miRNA predominantly from the 5p arm (≥90% of reads from the 5p arm); (ii) generating mature miRNA predominantly from the 3p arm (≥90% of reads from the 3p arm); and (iii) balanced (>10% of reads from each arm). As high-confidence miRNA genes, we considered genes coding for miRNA precursors annotated as “high confidence” in miRBase and/or deposited in MiRGeneDB v2.0. The precursors deposited in MiRGeneDB are defined based on criteria that include careful annotation of the mature versus passenger miRNA strands and evaluation of evolutionary hierarchy; therefore, they are much more credible than those in miRBase ^37,38^.

From the vcf.gz files, we extracted somatic mutation calls with PASS annotation. The extraction was performed with a set of in-house Python scripts (https://github.com/martynaut/pancancer_mirnome), an updated version of scripts used in our earlier research ^31^. To avoid duplicating mutations detected in multiple sequencing experiments in the same cancer patient, we combined files summing reads associated with particular mutations. Next, the lists of mutations detected by different algorithms were merged, removing multiple calls of the same mutations. To further increase the reliability of the identified mutations, we removed mutations that did not fulfill the following criteria: (i) at least two mutation-supporting reads in a tumor sample (if no mutation-supporting read was detected in the corresponding normal sample); (ii) at least 5× higher frequency of mutation-supporting reads in the tumor sample than in the corresponding normal sample; (iii) somatic score parameter (SSC) > 30 (for VarScan2 and SomaticSniper); and (iv) base quality (BQ) parameter for mutation-supporting reads in the tumor sample > 20 (for MuSE and MuTect2). We excluded hypermutated samples, defined as samples with > 10,000 mutations in the exome.

Target predictions for normal and mutant seed sequences were performed with the use of TargetScan Custom (release 4.2) ^39^, and secondary structure prediction was performed using mfold software (default parameters) ^40^. 3D pre-miRNA structures were predicted using RNAComposer software with default parameters ^41^ and visualized with PyMOL (Schrödinger, LLC, New York, NY, USA). Changes in motifs within miRNA precursors recognized by RNA-binding proteins were analyzed with a Python script based on miRNAmotif software ^42^.

### Statistics

Unless stated otherwise, all statistical analyses were performed with statistical functions from the Python module scipy.stats. Particular statistical tests are indicated in the text, and unless stated otherwise, a p-value < 0.05 was considered significant. If necessary, p-values were corrected for multiple tests with the Benjamini-Hochberg procedure.

Hotspot miRNA genes were identified based on the probability of occurrence of the observed number of mutations, which was calculated with the use of the 2-tailed binomial distribution, assuming a background random occurrence of identified mutations in all analyzed miRNA genes and considering the miRNA gene length. To further evaluate the reliability of the identified hotspot miRNA genes, we recalculated the mutation enrichment significance, weighting the mutation occurrences by the following factors: 2×, mutations in seeds (guide strand only); 1.5×, mutations in miRNAs (miRNA duplex), mutations affecting the functional motifs (identified by miRNAmotif) or +/-1 positions of DROSHA/DICER1 cleavage sites; and 1×, other mutations. Weight correction was not used to search for hotspot positions within miRNA genes.

For patient survival analyses, we used a log-rank test (from lifelines library ^43^) for specific cancers or a stratified version of the test for the Pan-Cancer cohort (survdiff function from statsmodels library ^44^). To determine the direction of mutation effects on survival, we used Cox’s proportional hazard model. Survival plots were created using KaplanMeierFitter from the lifelines library.

## Results

### Overview of miRNome mutations in TCGA cancers

To investigate the occurrence of somatic mutations in miRNA genes (miRNome), we took advantage of the WES datasets of 10,369 tumor/control sample pairs representing 33 different cancer types collected and analyzed by the TCGA project. The list of all cancer types and their abbreviations is provided in Table 1 [to avoid confusion, we will use the abbreviations only for the TCGA sample sets but not generally for particular types of cancer; in the latter case, we will use full cancer type names or alternative abbreviations indicated in the text]. We defined miRNome as 1918 miRNA genes (Supplementary Table 1) encompassing ~100 nt long fragments of genomic DNA coding for all pre-miRNAs (with 25 nt flanks) defined in miRBase v22.1, including 537 high-confidence pre-miRNAs annotated by miRBase and 466 knowledge-based expert-curated pre-miRNAs annotated in MirGeneDB v.2.0. It should be noted, however, that not all miRNA genes were covered by TCGA WES. In total, we found 10,588 mutations in miRNA genes; however, as the number of miRNome mutations in hypermutated samples (samples with >10,000 mutations in the whole exome) was highly correlated with the general mutation burden in these samples (Fig. 1a, Supplementary Fig. 1) and therefore is likely highly enriched in randomly occurring nonfunctional mutations, we decided to remove the hypermutated samples (114, ~1% samples; 3,478, ~33% mutations) from further analysis. The remaining 7,110 mutations (Supplementary Table 2), including 6,312 substitutions, 198 insertions, and 600 deletions, were found in 1,179 distinct miRNA genes. At the Pan-Cancer level, 3,370/10,255 (33%) samples had at least one miRNome mutation. This number was the highest in SKCM (298/460, 65%), DLBC (22/37, 59%), LUSC (285/497, 57%), and ESCA (99/181, 55%) and the lowest in PCPG (14/164, 9%), PRAD (39/497, 8%), and THCA (20/495, 4%) (Fig. 1c, Table 1). It should be noted, however, that the occurrence of mutations in miRNome is consistent with the general burden of mutations in particular cancer types (Fig. 1a,b, Supplementary Fig. 1). Additionally, as shown in Fig. 1d, there is a substantial fraction of samples with more than one mutation in miRNome. It is also noteworthy that some cancers, including COAD, STAD, and UCEC, have substantially heightened numbers of indel mutations (Table 1), which is consistent with known cancerous mechanisms, including DNA repair defects, associated with those cancers ^45–47^.

**Figure 1:**
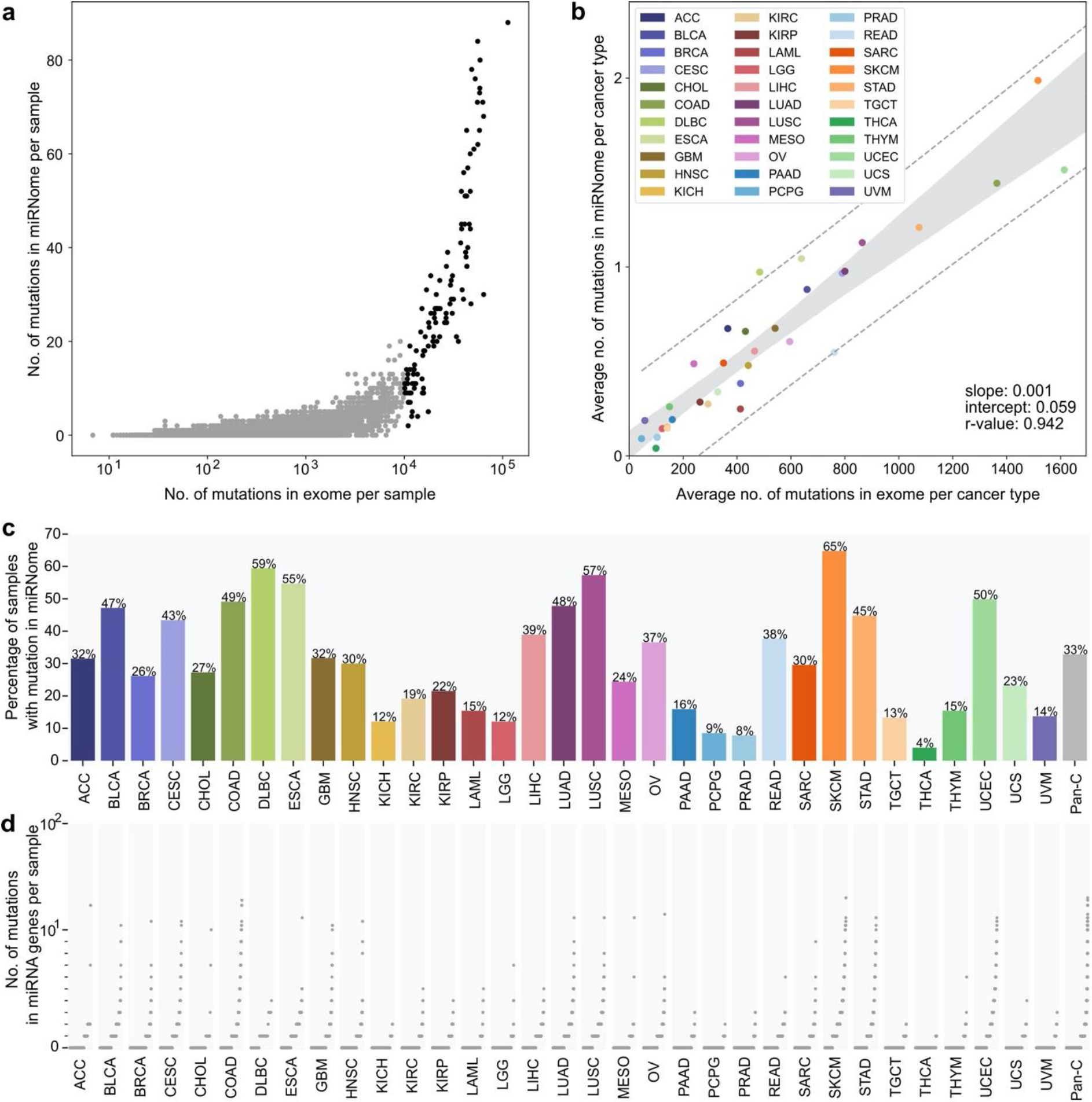
Summary of mutations identified in miRNome in TCGA cancer types. **(a)** The number of mutations in the exome (x-axis, log10 scale) and miRNome (y-axis, linear scale) per sample in Pan-Cancer. Each dot represents a single sample. Black dots represent hypermutated (>10k somatic mutations in exome) samples. **(b)** The average number of mutations in the exome (x-axis, linear scale) and miRNome per cancer type (y-axis, linear scale). Each dot represents a single cancer type. Hypermutated samples were excluded prior to the analysis. **(c)** Percentage of patients with at least one somatic mutation detected in miRNA genes. **(d)** The number of mutations in miRNA genes per patient. Each dot represents a single patient. Patients are ranked by the number of mutations in miRNA genes. Scale is linear (values 0-10) and log10 (values 10+).

**Table 1.**
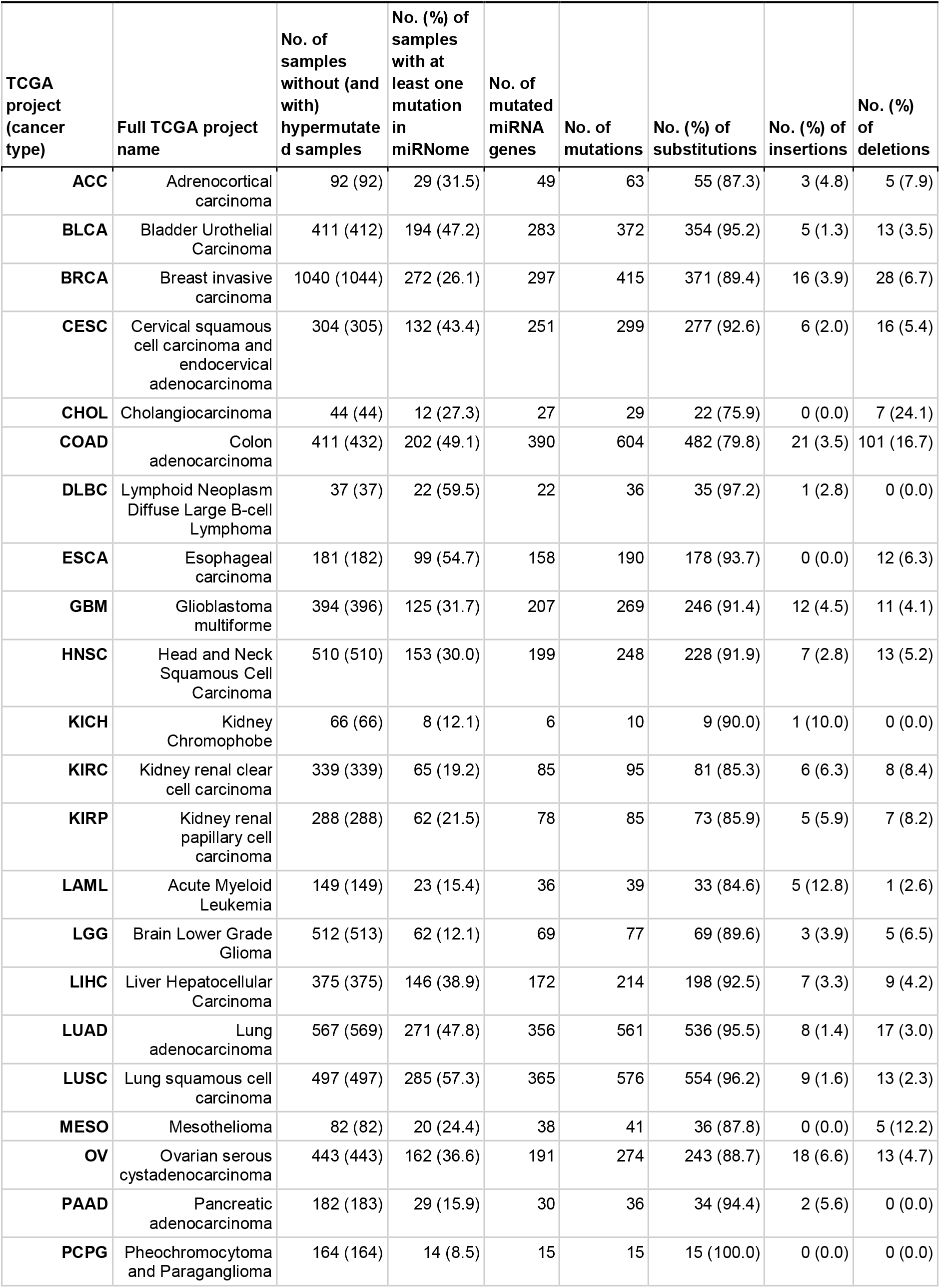
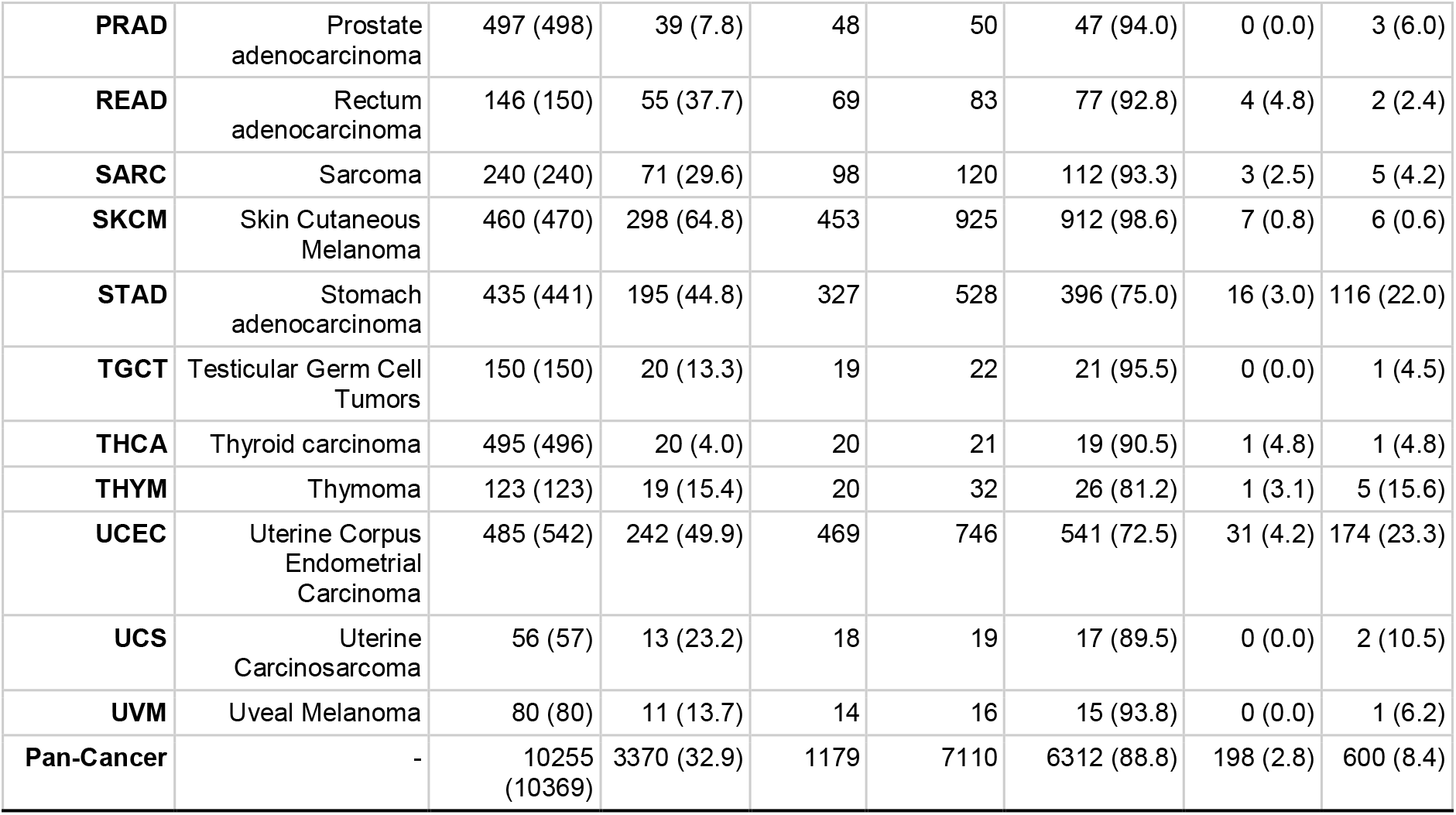
Summary of patients and mutations in miRNome per cancer type within the Pan-Cancer cohort.

### Localization of mutations within miRNA genes

For a closer examination of the localization of sequence variants in subregions of miRNA precursors, we superimposed the identified variants on the consensus miRNA precursor structure and categorized them according to localization in the miRNA gene subregions (Fig. 2a). The analysis shows that mutations occur in all regions of the miRNA gene, and in general, there is no strong imbalance in mutation localization within the miRNA precursor (Fig. 2b). A similar mutation distribution was observed when precursors of predominantly 5p- and 3p-miRNAs were analyzed separately (Fig. 2b, lower panels) and when the analysis was narrowed only to the high-confidence miRNA genes defined either by miRBase or miRGeneDB (Supplementary Fig. 2) or performed separately for individual cancer types (data not shown). We observed only a slightly decreased mutation rate in the 5p flanking region (Table 2); however, this effect may result from lower sequencing coverage in the flanking regions.

**Figure 2:**
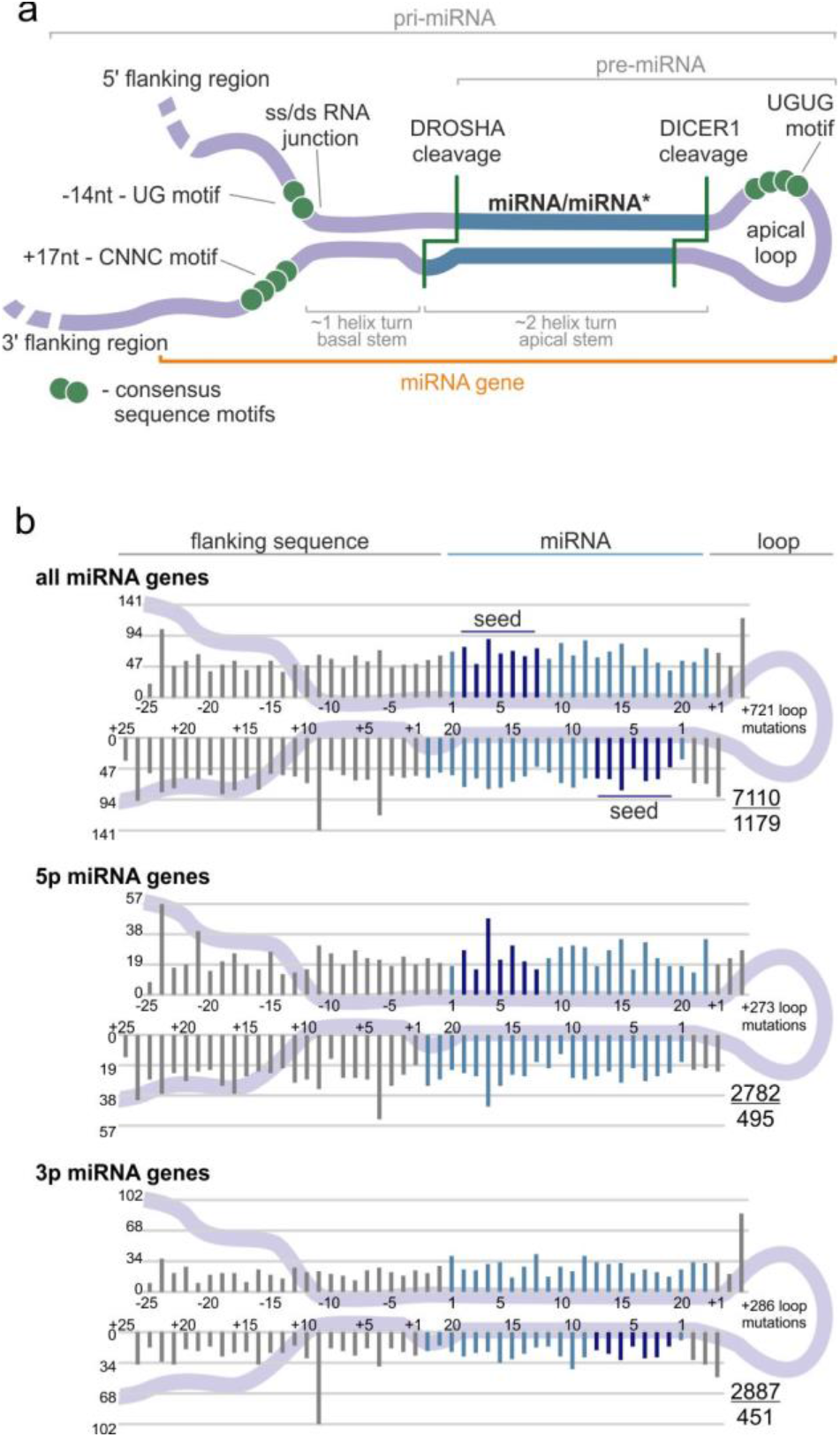
Localization of somatic mutations in miRNA precursors in Pan-Cancer. **(a)** An overview of a primary miRNA transcript with the indicated subregions considered in the study. The miRNA duplex is indicated in blue, and representative sequence consensus motifs recognized as enhancers of miRNA biogenesis are represented by green circles. **(b)** Localization of all detected mutations in the Pan-Cancer cohort. miRNA duplex positions are indicated in blue, seed regions in dark blue, and flanking regions and terminal positions of the apical loop in gray. The numbers in the lower-right corner represent the number of plotted mutations (upper) and the number of mutated miRNA genes (lower). If present, sequence variants localized beyond position 22 in longer mature miRNAs are shown cumulatively at position 22. The plot shows mutations within six positions of the loop (first 3 and last 3 nucleotides). The number of remaining mutations is indicated within the loop. Analyses were also performed in narrowed groups of miRNAs that release the guide miRNA strand predominantly from the 5p or 3p arm (lower panels).

**Table 2.**
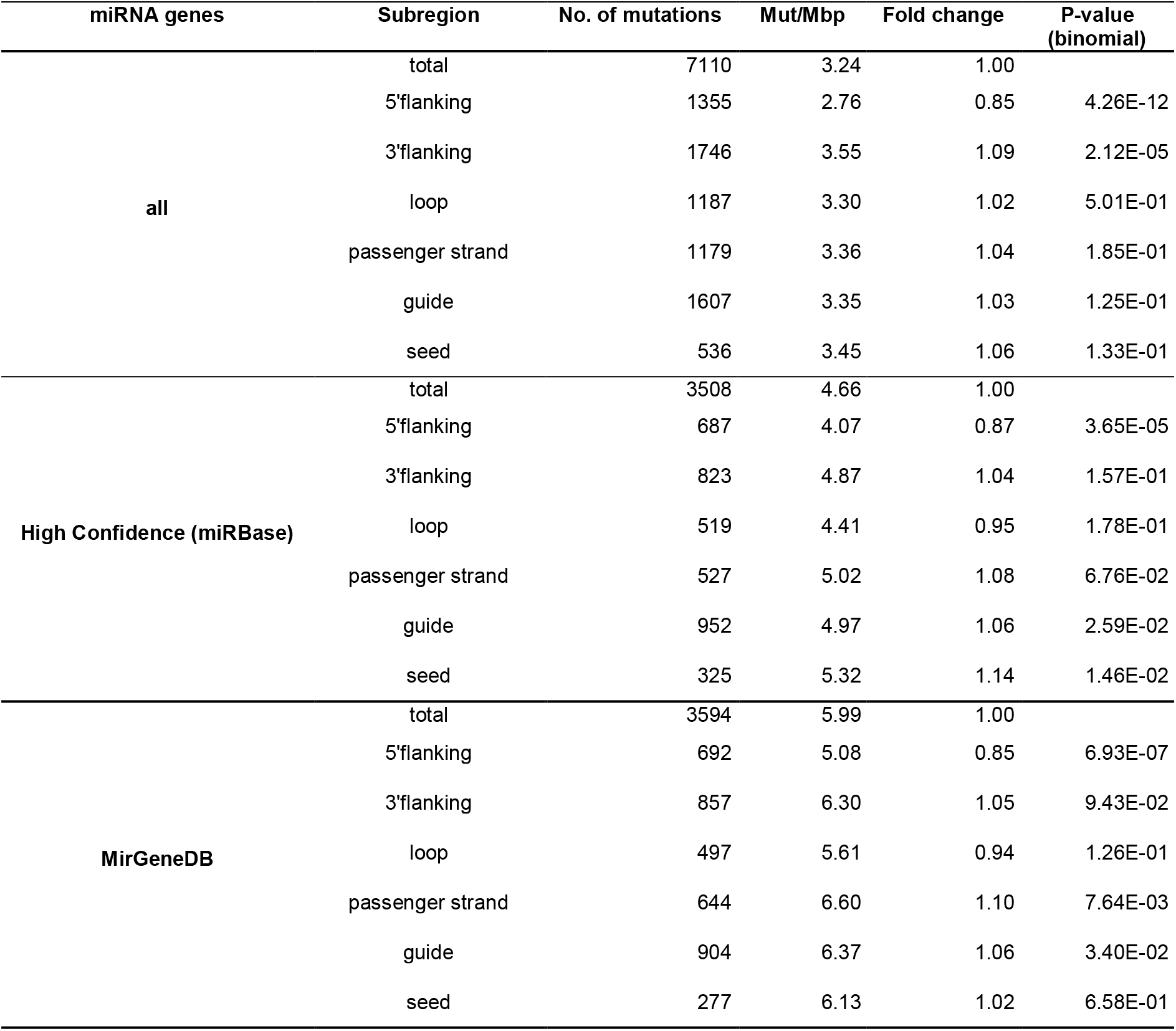
Distribution of somatic mutations within miRNA precursors.

| miRNA genes | Subregion | No. of mutations | Mut/Mbp | Fold change | P-value<br>(binomial) |
| --- | --- | --- | --- | --- | --- |
| **all** | total | 7110 | 3.24 | 1.00 |  |
|  | 5'flanking | 1355 | 2.76 | 0.85 | 4.26E-12 |
|  | 3'flanking | 1746 | 3.55 | 1.09 | 2.12E-05 |
|  | loop | 1187 | 3.30 | 1.02 | 5.01E-01 |
|  | passenger strand | 1179 | 3.36 | 1.04 | 1.85E-01 |
|  | guide | 1607 | 3.35 | 1.03 | 1.25E-01 |
|  | seed | 536 | 3.45 | 1.06 | 1.33E-01 |
| **High Confidence (miRBase)** | total | 3508 | 4.66 | 1.00 |  |
|  | 5'flanking | 687 | 4.07 | 0.87 | 3.65E-05 |
|  | 3'flanking | 823 | 4.87 | 1.04 | 1.57E-01 |
|  | loop | 519 | 4.41 | 0.95 | 1.78E-01 |
|  | passenger strand | 527 | 5.02 | 1.08 | 6.76E-02 |
|  | guide | 952 | 4.97 | 1.06 | 2.59E-02 |
|  | seed | 325 | 5.32 | 1.14 | 1.46E-02 |
| **MirGeneDB** | total | 3594 | 5.99 | 1.00 |  |
|  | 5'flanking | 692 | 5.08 | 0.85 | 6.93E-07 |
|  | 3'flanking | 857 | 6.30 | 1.05 | 9.43E-02 |
|  | loop | 497 | 5.61 | 0.94 | 1.26E-01 |
|  | passenger strand | 644 | 6.60 | 1.10 | 7.64E-03 |
|  | guide | 904 | 6.37 | 1.06 | 3.40E-02 |
|  | seed | 277 | 6.13 | 1.02 | 6.58E-01 |

### Significantly overmutated miRNA genes

In the next step, we searched for miRNA genes overburdened with mutations. The most frequently mutated miRNA genes are presented in Fig. 3a. We examined the numbers of mutations occurring in particular miRNA genes and the overall frequency of mutations in miRNome with the use of a binomial distribution test (p-value < 0.01) and showed that 81 of the recurrently mutated genes are significantly overmutated in Pan-Cancer (Table 3, Supplementary Table 3a). As mutations may not be randomly distributed in the genome and to consider the enrichment of functional variants, we next performed functionally weighted analysis, increasing the value of mutations located in most likely functional sequences including seed regions, DROSHA/DICER1 cleavage sites, miRNA duplexes, and protein binding motifs (see Materials and Methods section). The weighted analysis revealed 108 significantly overmutated miRNA genes, of which a substantial fraction overlapped with the genes identified in the ordinary binomial analysis (Table 3, Supplementary Table 3b). Two main types of overmutated miRNA genes can be distinguished based on mutation occurrence in various cancer types: one shows a somewhat sample-number dependent distribution of mutations across various cancer types (e.g., *hsa-miR-1324, hsa-miR-6891*, *hsa-miR-3675*), and the other shows an overrepresentation of mutations in one or two cancer types (e.g., *hsa-miR-1303* for STAD, *hsa-miR-890* for LUAD, *hsa-miR-519e* for OV).

**Figure 3:**
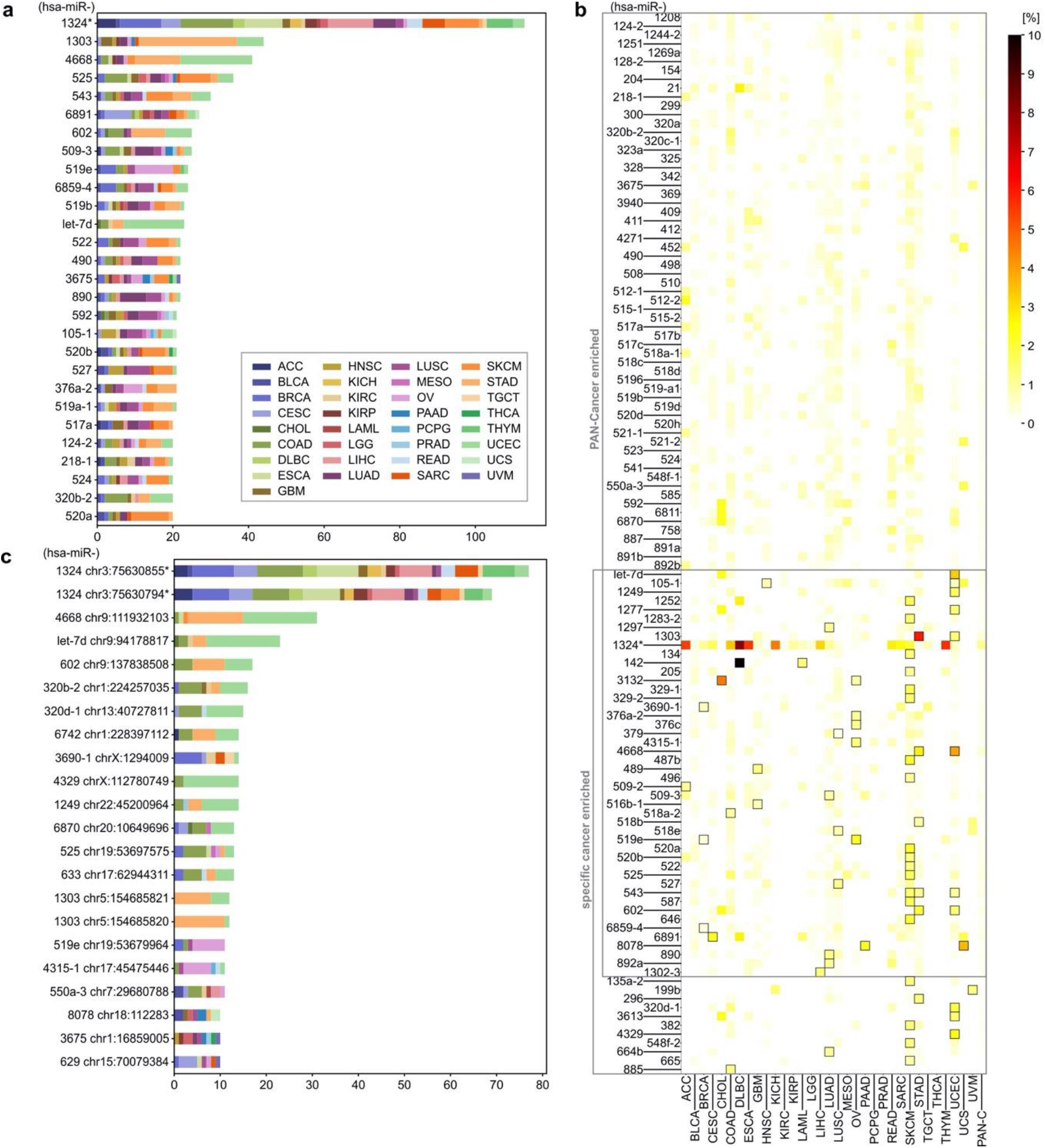
Most frequently mutated miRNA genes and hotspots in Pan-Cancer. **(a)** miRNA genes with at least 20 somatic mutations in Pan-Cancer. Each color represents a distinct cancer type. **(b)** Heatmap showing the percentage of mutations in miRNA genes overmutated (according to functionally weighted analysis) in Pan-Cancer and in specific cancers. Framed squares indicate cancer types in which gene enrichment reached statistical significance (adjusted p-value<0.01). Specific values of mutation frequencies are shown in Supplementary Table 4. **(c)** Hotspot positions in miRNA genes with at least 10 somatic mutations in Pan-Cancer (color legend as in panel **a**). To simplify the figure, we omitted the prefix hsa-miR in the gene IDs; * note the comment on mutations in hsa-miR-1324 at the end of the section *Examples of overmutated miRNA genes*.

**Table 3.**
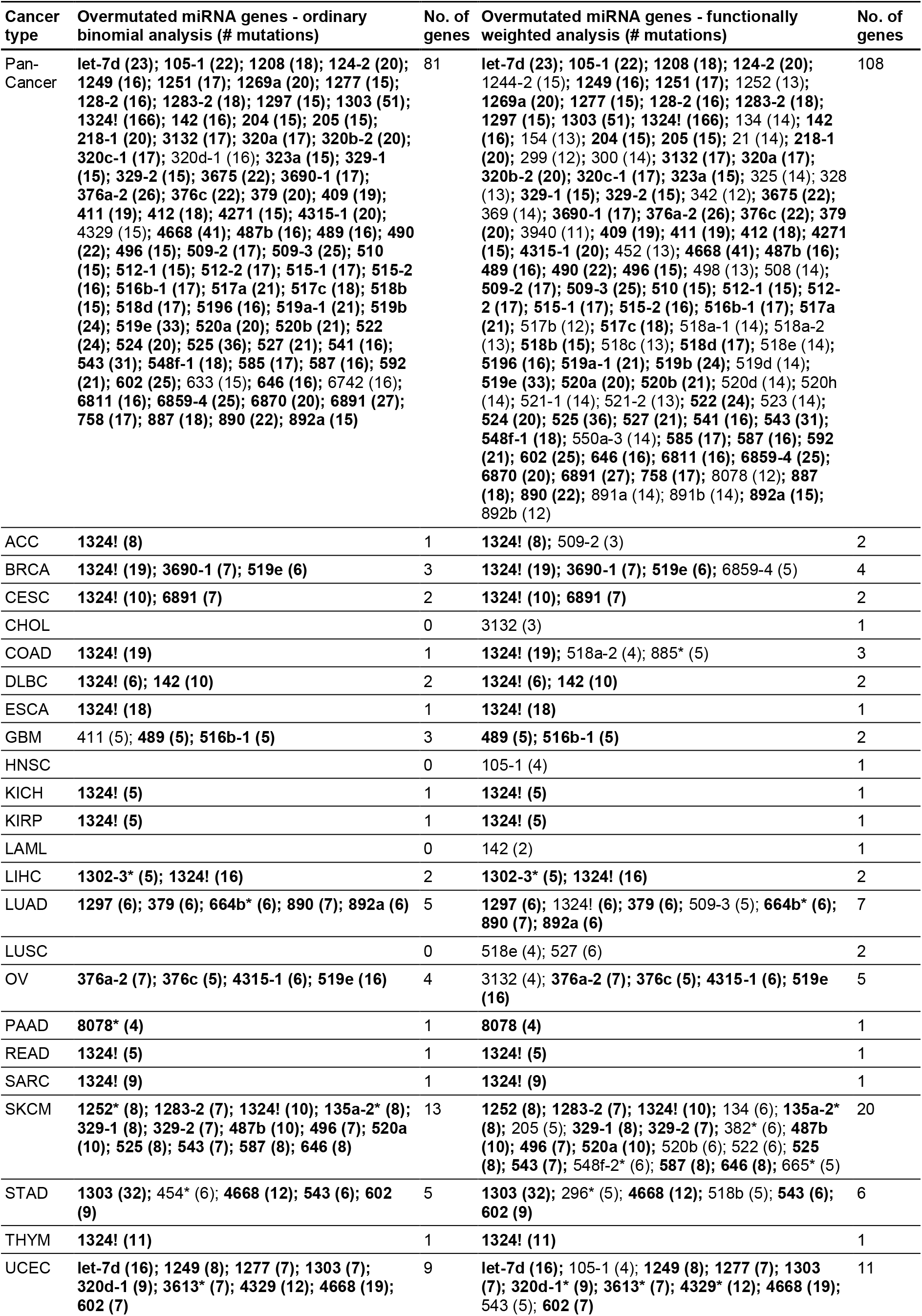
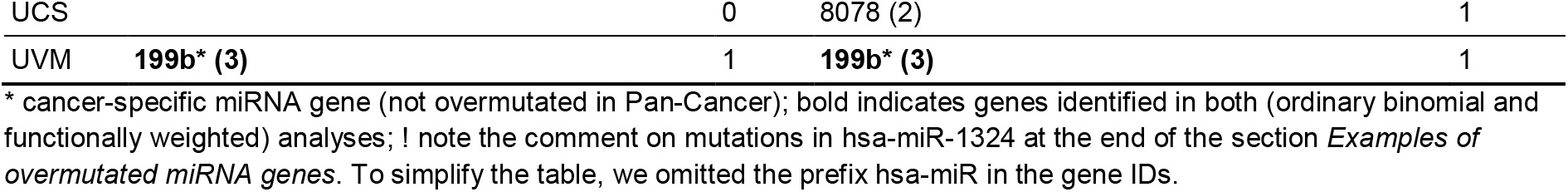
Summary of overmutated miRNA genes in Pan-Cancer and individual cancer types identified in ordinary binomial and functionally weighted mutation enrichment analysis.

As some mutations were unevenly distributed across cancer types, we also performed mutation enrichment analysis for individual cancers. This analysis revealed 55 and 80 additional miRNA genes overmutated in individual cancers in the ordinary binomial and functionally weighted analyses, respectively (Table 3, Supplementary Table 3). These lists included 8 and 12 cancer-specific overmutated genes, respectively, i.e., genes enriched in mutations in one or more cancers but not in Pan-Cancer. Among the most striking examples of the cancer-specific overmutated genes are (i) *hsa-miR-3613* with 7 mutations in UCEC but also with 1 mutation in CHOL and 1 in ESCA, (ii) *hsa-miR-135a-2* with 8 mutations in SKCM and 2 in UCEC, and (iii) *hsa-miR-664b* with 6 mutations in LUAD and 1 or 2 in UCEC, STAD, SKCM, and CESC. The highest overlap of overmutated miRNAs was observed between STAD and UCEC, which shared 4 out of 6 and 11 miRNA genes, respectively, according to functionally weighted analysis. The highest number of overmutated miRNA genes per cancer was found for SKCM (20) and UCEC (11). Some cancer types had no overmutated miRNA genes. As cancer type groups consisted of very different numbers of samples, in Fig. 3b and Supplementary Table 4, we visualized the occurrence of mutations in overmutated genes as a percentage of patients in the individual cancers and Pan-Cancer. As shown in Fig. 3b, the highest frequency of mutations belonged to *hsa-miR-142* in DLBC, but other overmutated genes often exceeded a frequency of 2% or even 5% in individual cancers, e.g., *hsa-miR-1303* in STAD (6%) and *hsa-miR-3132* in CHOL (4,5%). For reasons explained in the next section, we do not comment here on mutations in *hsa-miR-1324*. The localization of mutations in each of the overmutated genes is graphically illustrated in Supplementary Fig. 3.

The area of somatic mutations in miRNA genes is scarcely researched; therefore, it is difficult to compare our results directly to those of other studies. However, among the cancer-specific overmutated miRNA genes, we identified *hsa-miR-142*, in which somatic mutations were found before in several studies (for details see below). We also confirmed the recurrence of mutations in *hsa-miR-21* as previously identified with the Annotative Database of miRNA Elements (ADmiRe) ^48^. Not surprisingly, the current results overlap almost perfectly with our earlier results obtained for LUAD and LUSC ^31^. Minor discrepancies result from some differences in the technical approach (see Materials and Methods).

### Significantly overrepresented recurring point mutations

In the next step, we tested which recurrently mutated nucleotide residues are significant hotspots, i.e., positions mutated more frequently than expected by chance, taking into account overall mutation frequency and the number of samples in a particular cancer or the Pan-Cancer dataset. As such analysis may be strongly affected by the uneven occurrence of mutations in different genomic regions and different sequence contexts, to minimize false-positive results, we set a very stringent threshold of significance, an adjusted p-value<0.0001. The analysis showed 62 hotspots in Pan-Cancer and 69 in individual cancers, including 5 cancer-specific hotspots, 1 for DLBC, 2 for OV, and 2 for SKCM (Table 4, Supplementary Table 5). The two most frequently recurring point mutations were found in *hsa-miR-1324* (chr3:75630855T>C[+] and chr3:375630794C>G[+], Fig. 3c). Other interesting hotspot mutations include *hsa-miR-142* (chr17:58331260A>G[-] in the seed sequence of miR-142-3p) found in DLBC (3 mutations) and *hsa-miR-519e* (chr19:53679964G>A/T[+] and chr19:53679965G>A[+]) found in Pan-Cancer and OV. Please note that mutation occurring in a miRNA gene encoded on the minus chromosome strand in the sequence of a miRNA precursor occurs in reverse/complementary orientation; therefore, to avoid, confusion, a [+] or [-] sign indicates the orientation of the affected gene. As the recurrence of some mutations may be artifacts of not efficient filtering of germline variants, we checked the overlap of hotspot positions with the positions of SNPs. Although, due to the very large number of currently annotated SNPs, some of the SNPs coincide with the detected mutations, the very low population frequency of the SNPs or their type preclude confusing the SNPs with the recurrent mutations (Supplementary Table 5).

**Table 4.** Summary of hotspot positions found in miRNA genes in Pan-Cancer and individual cancer types identified in the binomial analysis.

| Cancer type | Significant hotspots (# mutations) | No. of hotspots |
| --- | --- | --- |
| Pan-Cancer | let-7d chr9:94178817 (23); 1208 chr8:128150186 (7); 1244-2 chr5:118974595 (7); 1249 chr22:45200964 (14); 1277 chrX:118386402 (7); 1302-3 chr2:113583038 (5); 1302-7 chr8:141786250 (5); 1303 chr5:154685809 (6); 1303 chr5:154685819 (8); 1303 chr5:154685820 (12); 1303 chr5:154685821 (12); 1324 chr3:75630794! (69); 1324 chr3:75630855! (77); 1324 chr3:75630860 (5); 15a chr13:50049206 (5); 205 chr1:209432167 (5); 296 chr20:58817670 (6); 320b-2 chr1:224257035 (16); 320c-1 chr18:21683580 (8); 320d-1 chr13:40727811 (15); 320e chr19:46709336 (5); 328 chr16:67202382 (6); 3652 chr12:103930488 (8); 3658 chr1:165907957 (7); 3663 chr10:117167687 (5); 3675 chr1:16858942 (5); 3675 chr1:16859005 (10); 3690-1 chrX:1294009 (14); 4271 chr3:49274155 (8); 4313 chr15:75762211 (5); 4315-1 chr17:45475446 (11); 4329 chrX:112780749 (14); 454 chr17:59137873 (8); 4668 chr9:111932103 (31); 487b chr14:101046508 (5); 489 chr7:93483953 (6); 512-1 chr19:53666774 (6); 512-2 chr19:53669258 (6); 519e chr19:53679964 (11); 519e chr19:53679965 (7); 520b chr19:53701276 (5); 525 chr19:53697575 (13); 525 chr19:53697594 (5); 543 chr14:101032009 (6); 548f-1 chr10:54607963 (6); 550a-3 chr7:29680788 (11); 567 chr3:112112876 (6); 570 chr3:195699434 (5); 602 chr9:137838508 (17); 624 chr14:31014652 (8); 629 chr15:70079384 (10); 633 chr17:62944311 (13); 6742 chr1:228397112 (14); 6811 chr2:237510924 (5); 6821 chr22:49962899 (5); 6859-4 chr16:17058 (9); 6859-4 chr16:17089 (6); 6870 chr20:10649696 (13); 6875 chr7:100868123 (5); 6891 chr6:31355224 (5); 6891 chr6:31355262 (6); 8078 chr18:112283 (10) | 62 |
| ACC | 1324 chr3:75630794! (4); 1324 chr3:75630855! (3) | 2 |
| BRCA | 1324 chr3:75630794! (8); 1324 chr3:75630855! (9); 3690-1 chrX:1294009 (6) | 3 |
| CESC | 1324 chr3:75630794! (5); 1324 chr3:75630855! (5); 629 chr15:70079384 (4); 6891 chr6:31355224 (5) | 4 |
| COAD | 1324 chr3:75630794! (8); 1324 chr3:75630855! (10); 320b-2 chr1:224257035 (5); 320c-1 chr18:21683580 (4); 320d-1 chr13:40727811 (5); 525 chr19:53697575 (5); 602 chr9:137838508 (4); 633 chr17:62944311 (4) | 8 |
| DLBC | 1324 chr3:75630794! (3); 1324 chr3:75630855! (3); 142 chr17:58331260* (3) | 3 |
| ESCA | 1324 chr3:75630794! (8); 1324 chr3:75630855! (9); 624 chr14:31014652 (3) | 3 |
| KICH | 1324 chr3:75630855! (3) | 1 |
| KIRP | 1324 chr3:75630794! (3) | 1 |
| LIHC | 1302-3 chr2:113583038 (5); 1324 chr3:75630794! (7); 1324 chr3:75630855! (7) | 3 |
| OV | 1244-2 chr5:118974595 (3); 376a-2 chr14:101040083* (3); 376c chr14:101039694* (3); 4315-1 chr17:45475446 (6); 519e chr19:53679964 (7); 519e chr19:53679965 (4) | 6 |
| READ | 1324 chr3:75630855! (3) | 1 |
| SARC | 1324 chr3:75630794! (3); 1324 chr3:75630855! (5) | 2 |
| SKCM | 1324 chr3:75630794! (4); 329-1 chr14:101026832* (4); 487b chr14:101046508 (5); 524 chr19:53711049* (4); 525 chr19:53697594 (5) | 5 |
| STAD | 1208 chr8:128150186 (4); 1303 chr5:154685809 (4); 1303 chr5:154685819 (8); 1303 chr5:154685820 (11); 1303 chr5:154685821 (8); 296 chr20:58817670 (5); 454 chr17:59137873 (6); 4668 chr9:111932103 (12); 602 chr9:137838508 (7); 6742 chr1:228397112 (5) | 10 |
| THYM | 1324 chr3:75630794! (4); 1324 chr3:75630855! (7) | 2 |
| UCEC | let-7d chr9:94178817 (16); 1249 chr22:45200964 (8); 1277 chrX:118386402 (7); 1303 chr5:154685821 (4); 320b-2 chr1:224257035 (6); 320c-1 chr18:21683580 (4); 320d-1 chr13:40727811 (8); 3658 chr1:165907957 (5); 4271 chr3:49274155 (6); 4329 chrX:112780749 (12); 4668 chr9:111932103 (16); 602 chr9:137838508 (6); 633 chr17:62944311 (4); 6742 chr1:228397112 (5); 6870 chr20:10649696 (5) | 15 |

\* cancer-specific hotspots (not overmutated in Pan-Cancer); ! note the comment on mutations in hsa-miR-1324 at the end of the section *Examples of overmutated miRNA genes*. To simplify the table, we omitted the prefix hsa-miR in the gene IDs.

### Examples of overmutated miRNA genes

#### Hsa-miR-142

*Hsa-miR-142* is, to the best of our knowledge, the only miRNA gene convincingly shown to be recurrently mutated in several neoplasms, which include acute myeloid leukemia (AML) ^49,50^ and different types of B-cell lymphoma ^51,52^, chronic lymphocytic leukemia (CLL) ^53^ and diffuse large-cell B-cell lymphoma ^54–57^. Additionally, in our sequencing analysis performed within the framework of other projects, we found also one mutation in the seed region of *hsa-miR-142* (chr17:58331263C>T[-]) in the Raji Burkitt lymphoma cell line (out of 5 Burkitt’s lymphoma cell lines tested) (Fig. 4a). The occurrence of *hsa-miR-142* mutations in hematological cancers may be consistent with the high abundance of miR-142-3p in mature hematologic cells ^58,59^ and with the observation that loss of the miRNA impairs the development and function of different hematologic lineages ^60–62^.

**Figure 4:**
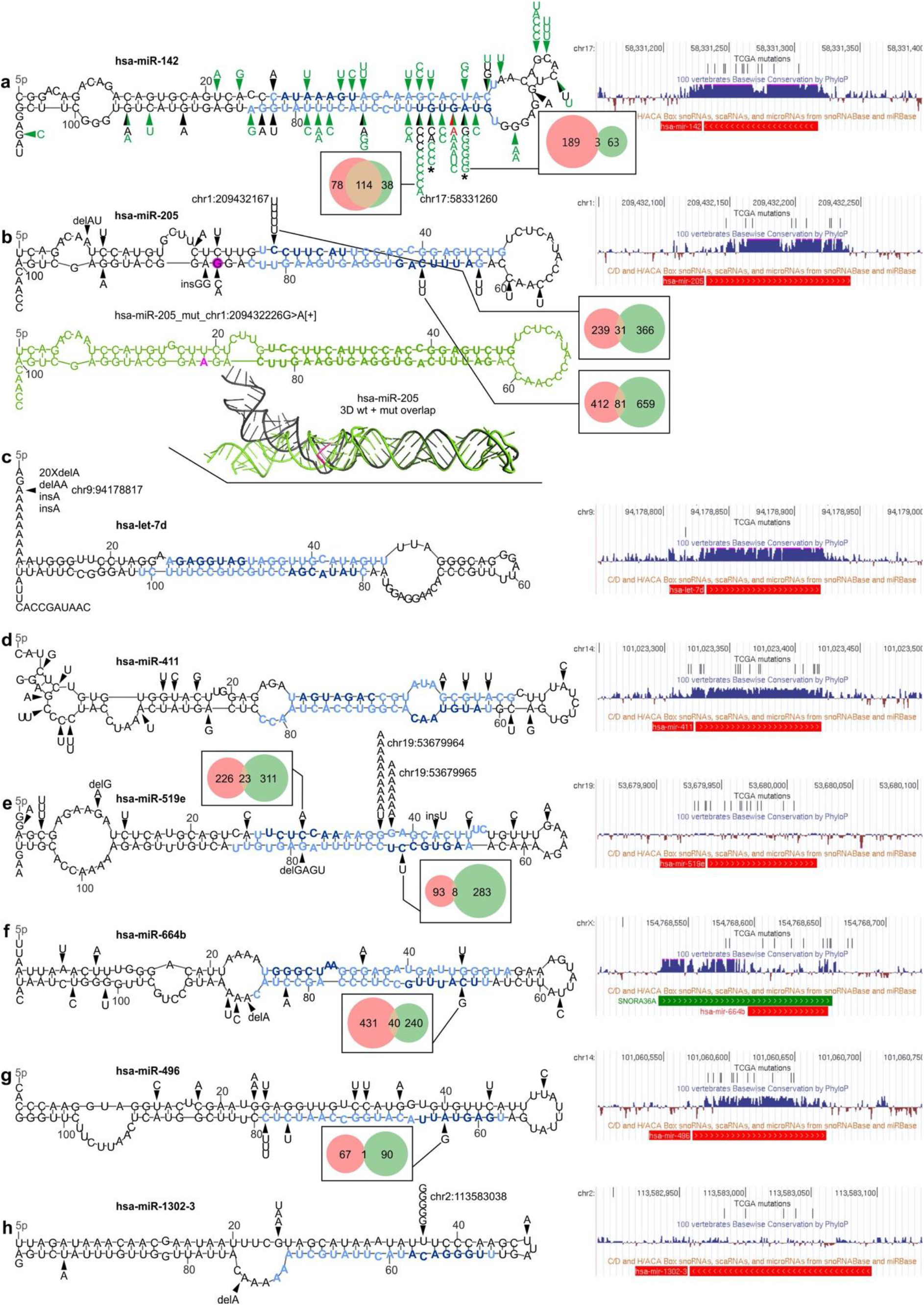
Localization of mutations in the selected overmutated miRNA genes. In each panel, on the left, mutations identified in the study in the TCGA samples (black arrowheads) are shown on mfold-predicted 2D structures of the miRNA precursors. Mature miRNA and seed sequences are indicated in light blue and dark blue, respectively. On the right, a screenshots from the UCSC genome browser, showing (from the top) a custom track with positions of the TCGA mutations detected in the study, conservation of the genomic region, and position of short noncoding RNAs, including pre-miRNAs (red bar; according to miRbase) and snoRNAs (green bar). For selected seed mutations, Venn-diagrams indicate the number of predicted targets for the wild-type (pink circle) and mutated (green circle) seeds. Genomic positions of the most significant hotspots are indicated next to mutations symbol. (a) Mutations found in *hsa-miR-142*. Green arrowheads indicate mutations detected in previous studies ^49–53,55–57^ (for details see Supplementary Table 6); red arrowhead indicates the mutation identified by us in the Burkitt’s lymphoma (Raji) cell line. * indicates the mutations tested functionally ^27^. (b) Mutations found in *hsa-miR-205* (above). Below: (i) the corresponding 2D structure of the precursor with the chr1:209432226G>A[+] mutation (drawn in green) and (ii) superimposed 3D structures of the wild-type (black) and mutant (green) precursors are shown. The position of the mutation and the mutant allele are marked in pink. (c-h) Mutations found in *hsa-let-7d, hsa-miR-411, hsa-miR-519e, hsa-miR-664b, hsa-miR-496*, and *hsa-miR-1302-3*, respectively.

In this study, we found 16 mutations in *hsa-miR-142*. Consistent with previous studies, we identified the highest number of mutations in DLBC (10 mutations, including 3 in one sample) and LAML (2 mutations), but we also found 4 mutations in solid tumors, i.e., in UCEC, BLCA, GBM, and BRCA, in which the *hsa-miR-142* have not been found before. Five of the mutations in DLBC are located in the seed sequence of miR-142-3p, three in the 7th nucleotide (significantly recurring position chr17:58331260A>G[-]) and two in the 6th nucleotide (chr17:58331261C>G/T[-]) of the seed. Additionally, one mutation (chr17:58331264T>C[-]) was detected in the 3rd seed nucleotide in LAML. To better understand the distribution of mutations in *hsa-miR-142*, we combined the mutations detected in our study with the mutations detected previously (Fig. 4a, Supplementary Table 6). The distribution of mutations shows pronounced clustering of the mutations in the miR-142-3p seed region, with chr17:58331260A[-] being the most frequently mutated nucleotide, substituted with either G[-] (n=8) or T[-] (n=1). Nonetheless, a substantial fraction of the mutations is dispersed in other parts of the gene, including two recurring mutations in two subsequent positions of the loop. This result may suggest that the miRNA hairpin precursor is quite a fragile structure, and therefore, almost any mutation may be deleterious for the gene, either by disturbing the structure of the precursor or by disruption of the seed sequence. A recent functional study of two seed mutations, i.e., chr17:58331264T>C[-] and chr17:58331261C>G[-], showed that even though the mutations are located in miR-142-3p, they result in a decrease in both miR-142-3p and miR-142-5p levels and reverse the miR-5p:3p ratio (in favor of miR-3p) ^27^. The functional consequences of the mutations are (i) aberration of hematopoietic differentiation, enhancing the myeloid and suppressing the lymphoid potential of hematopoietic progenitors, and (ii) inefficient repression of *ASH1L*, resulting in increased levels of HOXA9 and A10 (positively regulated by ASH1L) and ultimately leukemic transformation ^27^. Although the effects of other mutations were not directly tested, it is likely that they are also deleterious mutations, resulting in similar functional consequences to the two functionally validated mutations. Otherwise, it would be difficult to explain their recurrence in specific cancers, especially in relatively low-mutation hematologic neoplasms. The question remains whether mutations in solid tumors, in which miR-142-3p acts predominantly as a tumor suppressor, among others targeting and downregulating *TGFB1R* and *HMGB1* ^63,64^, may also have functional consequences. It was shown that miR-142-3p plays a role in different solid tumors (e.g., breast ^65,66^, ovarian ^67^, colorectal ^68,69^, and lung cancer ^70–72^). *Hsa-miR-142* constitutes an example showing that cancer-specific enrichment of mutations may be a strong indicator of their functional relevance for cancer.

#### Hsa-miR-205

Among the highly mutated miRNA genes, several encode miRNAs with an important and well-documented role in cancer. These genes include *hsa-miR-205*, whose miR-205-5p acts predominantly as a suppressormiR but also, depending on a tumor context and/or expression profile, as an oncomiR (reviewed in ^73^), among others in breast, prostate, and lung cancer. miR-205 is a highly conserved and well-validated miRNA. We found that *hsa-miR-205* was overmutated in Pan-Cancer (in total, 15 mutations) with mutations in SKCM (5 mutations), CESC (3), LUSC (2), BLCA (2), COAD, ESCA, and THYM. Five of the mutations are located in a single hotspot position (chr1:209432167C>T[+]) that is the first position of the seed sequence of miR-205-5p (guide miRNA). As we have shown before, the mutation may substantially affect target recognition, disrupting 250/288 (87%) predicted miR-205-5p targets and creating 471 new targets ^31^. The disrupted targets include many validated miR-205-5p targets, including the oncogenes *VEGFA* (mediator of angiogenesis) and *E2F1* (transcription factor controlling cell cycle), whose less effective downregulation may trigger tumor progression, invasion and/or metastasis ^74,75^. The other disrupted validated targets include *MED1, ERBB3*, and *PTEN* ^73^. On the other hand, the recurrence of a specific mutation may suggest its gain-of-function character, such as the creation of a new seed/miRNA targeting the gene, whose downregulation may be beneficial for cancer. Examples of predicted targets (TargetScan) of mutated miR-205-5p include proapoptotic *TP53BP2* ^76^, suppressor gene *ARID1A* ^77^, and *DLC1* ^78^. The other *hsa-miR-205* mutations are dispersed alongside the miRNA duplex, hairpin loop, and flanking sequences. The mutations may affect the structure of the miRNA precursor and consequently its processing and effective miRNA biogenesis. An example of a mutation seriously affecting the structure is chr1:209432226G>A[+] transition, which disrupts the structure in the DROSHA cleavage site (Fig. 4b).

#### Hsa-let-7d

Another highly mutated miRNA gene playing a role in cancer is *hsa-let-7d*, which is located in a *let-7a-1/let-7f-1/let-7d* cluster and belonging to the let-7 family, one of the most extensively studied miRNA families in cancer. The let-7 miRNAs were found to be downregulated in many cancers. They act as suppressormiRs, among other roles, directly targeting *RAS* oncogenes, *HMGA2*, and other genes playing a key role in processes such as the cell cycle, proliferation, and apoptosis ^79,80^. More specifically, it was shown that let-7d targets genes such as *KRAS, LIN28*, and *MYC* (reviewed in ^81^). *Hsa-let-7d* is a highly conserved and well-validated miRNA gene. We found that *hsa-let-7d* was overmutated in Pan-Cancer (in total 23 mutations) and UCEC (16 mutations) but was also recurrently mutated in STAD (3) and COAD (2). All the mutations are indels of the poly-A10 tract (chr9:94178817[+] delA (20), delAA (1), and insA (2)) located in a 5p flanking sequence of the let-7d precursor (Fig. 4c). Although the mutation does not directly affect the sequence of mature let-7d, it may still affect miRNA processing. It should be noted, however, that indels in the polynucleotide tract, such as those observed in *hsa-let-7d*, may result from microsatellite instability (MSI), which often occurs in different cancers associated with impairment of DNA mismatch repair mechanisms. This is consistent with the overrepresentation of the *hsa-let-7d* indels in cancers such as UCEC, STAD, and COAD, in which MSI is especially frequent ^47^. On the other hand, it was suggested that MSI-associated mutations may constitute genuine functional variants ^82^.

#### Hsa-miR-411

*Hsa-miR-411* is overmutated in Pan-Cancer (19 mutations) and GBM (5 mutations) and is also recurrently mutated in SKCM (3) and ESCA (2). All mutations in hsa-miR-411 are substitutions and are generally dispersed over the gene without clustering in any specific region or hotspot (Fig. 4d), resembling a pattern of loss-of-function mutations, usually characteristic of tumor suppressor genes. This finding may be consistent with a predominantly tumor suppressor role and downregulation of miR-411-5p and miR-411-3p in different cancers, including glioblastoma, breast cancer, ovarian cancer, lung adenocarcinoma, bladder cancer and renal cell carcinoma ^83–87^. For example, miR-411-5p suppresses cell growth, migration, and invasion, and its downregulation correlates with lymph node metastasis in breast cancer ^84^ and contributes to chemoresistance in ovarian cancer ^88^. Consistent with the potential function of the *hsa-miR-411* mutation in GBM, miR-411-5p is considered a brain-enriched miRNA and was previously associated with neurological and neuropsychiatric disorders ^89,90^. Interestingly, miR-411-5p is posttranscriptionally modified (substitution A>I (inosine) in the 5th position of miR-411-5p seed) in both normal brain and glioblastoma multiforme tissues ^87,91^. The precise function of this modification is not known; however, *in vitro* analyses revealed that pri-miRNA editing is likely to interfere with miRNA processing ^92^. Additionally, as modification occurs in the seed sequence, it may also influence the recognition of miRNA targets ^92^. It is likely that, at least in some cases, although the observed mutations do not correspond precisely with the position of the A>I modification, the mutations may mimic the transient effect of the modification. Mutation-induced imprecise processing of the precursor may also influence 5p/3p strand dominance and the generation of 5p heterogeneity of miR-411-5p, which were shown to differentiate normal brain and glioblastoma tissues ^87^.

#### Hsa-miR-519e

With 33 identified mutations, *hsa-miR-519e* is overmutated in Pan-Cancer and two female-specific cancers, OV (16) and BRCA (6). The majority of the mutations are located in two subsequent hotspot positions, i.e., the 12th and 13th nucleotides (chr19:53679964G>A/T[+] and chr19:53679965G>A[+]) of the miR-519e-5p (passenger) strand (Fig. 4e), which are mutated predominantly in OV. *Hsa-miR-519e* belongs to the large (>50) miR-515 family coded in a cluster on 19q13.42. It is not a highly validated miRNA gene and is not well recognized as a cancer-related miRNA; however, it was shown to have decreased levels in epithelial ovarian cancer, increased levels in ovarian cancer ^93^, and decreased levels upon 17ß-estradiol (E2) treatment in the MCF-7 breast cancer cell line ^94^ and was shown in a cell line experiment to influence anticancer drug chemosensitivity ^95^.

#### Hsa-miR-664b

*Hsa-miR-664b* is significantly overmutated in LUAD, with 6 mutations found in this cancer and 6 in other cancer types, including UCEC, STAD, SKCM, and CESC. The mutations were dispersed throughout the entire gene, with 5 mutations located in the mature miRNA sequences (Fig. 4f). *Hsa-miR-664b* is a highly validated (miRBase) and moderately conserved miRNA gene that substantially overlaps with the *SNORA36A* H/ACA box small nuclear RNA (snoRNA; snoRNABase) playing a role in the pseudouridylation of rRNAs and snRNAs; therefore, all mutations may also affect the function of snoRNA. In our previous study, we showed that some of the mutations, especially chrX:154768615G>T[+] and chrX:154768652C>A[+], cause serious structural aberrations ^31^. miR-664b-5p was also recently shown to act as a cancer suppressor in hepatocellular cancer cell lines ^96^. The downregulated miR-664b-5p was associated with lower overall survival in cervical cancer ^97^, the proliferation of cutaneous malignant melanoma cells ^98^, and the progression of breast cancer ^99^. The role of the miRNA is associated with the downregulation of *AKT2* ^96^.

#### Hsa-miR-496

*Hsa-miR-496* was overmutated in the Pan-Cancer cohort with 15 mutations. Additionally, it was overmutated in SKCM with 7 mutations, 3 of which were located at a single position (chr14:101060649C>T[+]) in the DROSHA cleavage site (Fig. 4g), which may affect both the efficiency and precision of miRNA excision. Other mutated cancers include LUSC (2 mutations), HNC (2 mutations) and OV, UCEC, LUAD, and GBM (with single mutations). *Hsa-miR-496* is a conserved miRNA gene located in a large cluster (~40 miRNA genes) at ch14q31.31. It was shown that miR-496-3p plays a role in the regulation of the mTOR pathway ^100^ and Wnt pathway-mediated tumor metastasis in colorectal cancer ^101^.

#### Hsa-miR-1302-3

*Hsa-miR-1302-3* is an example of a cancer-specific overmutated miRNA gene that is overmutated only in LIHC (5 mutations). Other mutated cancers include STAD (2 mutations) and ESCA, LUSC, CESC, and BRCA (with single mutations). Most of the mutations occur in two positions in the 5’ arm of the precursor, one of which is a hotspot mutation (chr2:113583038A>C[-]) significantly recurrent in LIHC and PAN-Cancer, localized within the passenger miRNA strand (Fig. 4h). The mutation replaces a U with G in a U:C mismatch and thus replaces the mismatch with a Watson-Crick pair G:C, greatly stabilizing the hairpin structure of the precursor (ddG=-6 kcal/mol; RNA mfold). miR-1302-3p has not been broadly researched, but its upregulation is associated with the recurrence and metastasis of prostate cancer ^102^. It is also connected with infertility and potentially with breast cancer through the regulation of the *CGA* gene ^103,104^.

#### Hsa-miR-1324

Finally, *hsa-miR-1324* is the most commonly mutated gene, with a total of 166 mutations in Pan-Cancer, greatly exceeding the other highly mutated genes. The vast majority of the mutations (n=146) were located in just two positions (i.e., chr3:75630855T>C[+] and chr3:375630794C>G[+]), which are also the two most highly mutated hotspots (Supplementary Fig. 3). We observed similar high frequency and a similar pattern of mutations in *hsa-miR-1324* in a relatively small panel of diffuse large B-cell lymphoma and Hodgkin lymphoma cell lines sequenced in our laboratory with the conventional Sanger sequencing method (data not shown). However, the analysis of the genomic location of *hsa-miR-1324* revealed that it is embedded in a large (>10 kb) segmentally duplicated region highly similar (>>95%) to at least 4 other sequences in the genome and likely variable in copy number ^29,105,106^. The detailed comparison of the *hsa-miR-1324* sequence with its paralog counterparts revealed that the substitutions differentiating paralogs correspond (position and type of substitution) with the identified mutations. This indicates that the *hsa-miR-1324* mutations are most likely artifacts of the sequencing procedures and/or computational analyses (e.g., mapping). The additional facts arguing against the genuineness of the *hsa-miR-1324* mutations are (a) relatively low fraction of an alternative allele, (ii) relatively high presence of the mutated reads in the reference noncancerous samples, (iii) frequent identification of the mutations by only one mutation caller algorithm, and (iv) observation of similar sequence anomalies in our own NGS experiments both in cancer and noncancerous samples. Additionally, miR-1324 is a low-confidence miRNA with only 48 confirming reads (miRBase, Mar 17, 2020) and is not annotated in MirGeneDB. In summary, based on the above facts, we concluded that the *hsa-miR-1324* alterations are not credible somatic mutations, and therefore, we did not pursue further analysis of miR-1324.

### Effect of the mutations on the expression of the affected miRNA genes

To check whether mutations may affect miRNA expression, we compared the levels of miRNAs in samples with mutations vs. samples without mutations in genes either overmutated or with hotspot mutations in a particular cancer type or Pan-Cancer. To level the between-cancer expression differences, prior to Pan-Cancer analysis, we normalized the level of each miRNA to make its median level (equal to 0) and variation comparable between cancer types. We took into account only miRNAs whose level was >0 in at least 70% of the analyzed samples. Notably, not all miRNAs were covered in the TCGA miRNA expression data. As a result of the analysis, we identified 10 miRNA genes whose miRNA levels were downregulated in mutated samples, including *hsa-miR-134*, for which both miR-134-5p and miR-134-3p were downregulated in Pan-Cancer (Supplementary Table 7, and Fig. 5a). Additionally, we found 2 miRNAs whose levels were downregulated by particular hotspot mutations (Fig. 5a). No miRNA was upregulated by the mutations. The striking excess of downregulated miRNAs is consistent with the notion that most mutations are loss-of-function mutations for particular miRNA genes. It should be noted, however, that due to a low number of mutations, especially in the hotspots, the analysis is of relatively low statistical power, and most results are only nominally significant (p<0.05; Supplementary Table 7).

**Figure 5:**
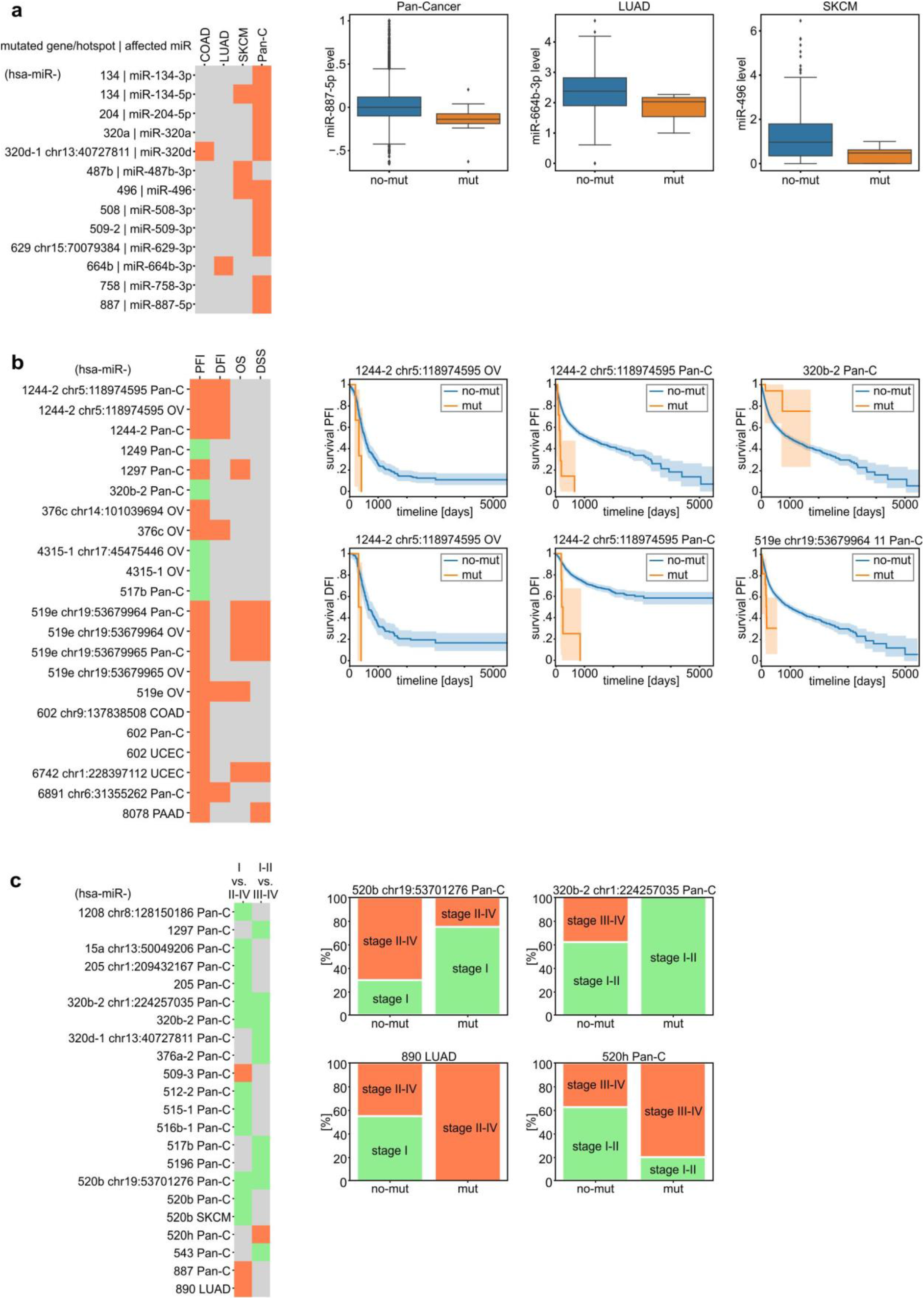
Association of miRNA gene mutations with miRNA expression, patient survival, and cancer stage. **(a)** Heatmap (on the left) shows miRNAs whose levels are significantly changed in samples mutated in the indicated genes or hotspots. Orange indicates that all miRNAs were downregulated by mutations in their genes. Box plots (on the right) show representative examples of miRNAs whose levels were changed in samples with mutations (mut) vs. samples without mutations (no-mut) in the corresponding gene. For Pan-Cancer analysis, miRNA levels were normalized to allow comparison between cancer types. **(b)** Heatmap (on the left) shows the miRNA genes or hotspot mutations significantly associated with PFI (green - positively, red - negatively). In the next columns, associations of the genes/hotspots with the other survival metrics (DFI, DSS, and OS) are also shown. Example survival plots comparing mut and no-mut samples are shown on the right. **(c)** Heatmap (on the left) shows miRNA genes or hotspot mutations associated with cancer stages. Red and green colors indicate associations of samples bearing mutations with higher and lower cancer stages, respectively. Examples of associations of mutation with the distribution of cancer stages are shown on the right. To simplify the figure, we omitted the prefix hsa-miR in the gene IDs.

### Association of mutations in miRNA genes with patient survival and cancer aggressiveness

Changes in miRNA expression, processing, and target specificity may influence various cancer-related processes, including cell proliferation, metastasis, progression, and/or drug resistance. These changes may result in disease progression and treatment outcomes affecting patient survival. Multiple metrics associated with patient survival have been gathered within the TCGA project, including overall survival (OS), disease-specific survival (DSS), disease-free interval (DFI), and progression-free interval (PFI), although not all are optimal for all cancers ^35^.

According to the recommendations in Liu et al. ^35^, we used PFI as a metric because it was permissible and most informative (had the highest statistical power) for the majority of TCGA cancer types. As survival metrics, including PFI, differ substantially between cancers, Pan-Cancer comparisons of survival in patients with mutations vs. patients without mutations may be affected by the fact that mutations are not equally distributed between cancer types. To overcome this effect, we used a stratified version of the log-rank test. We found 22 significant associations between mutations in the overmutated miRNA genes or hotspot positions and the PFI of cancer patients (either specific cancers or PAN-Cancer). The associations were linked with mutations in 12 distinct miRNA genes (Fig. 5b). Interesting examples may be (i) *hsa-miR-1244-2*, in which hotspot mutations chr5:118974595C>T[+] are associated with decreased PFI in both OV and Pan-Cancer and total mutations decrease PFI in Pan-Cancer; (ii) *hsa-miR-519e*, in which total mutations are associated with decreased PFI in OV and hotspot mutations (chr19:53679964G>A/T[+], chr19:53679965G>A[+]) decrease PFI in OV and Pan-Cancer; and (iii) *hsa-miR-602*, in which hotspot mutations (chr9:137838508GC>G[+]) decrease PFI in COAD and total mutations decrease PFI in Pan-Cancer and UCEC (Fig. 5b). Additionally, we observed that mutations in *hsa-miR-411* that was overmutated only in ordinary binomial analysis in GBM were associated with a decrease in PFI in GBM (p-value <0.001, data not shown). As shown in the left panel of Fig. 5b, mutations of particular miRNA genes associated with PFI are also frequently associated with other measures of survival (DFI, OS, DSS), which were analyzed as appropriate for particular cancers ^35^.

A profound excess mutations associated with decreased survival may suggest a predominant tumor suppressor role of miRNAs, which is also consistent with the global decrease in miRNA levels observed in many cancers.

Next, we compared the occurrence of mutations in miRNA genes with cancer stages. The analysis showed 25 statistically significant associations of mutations, predominantly with lower cancer stages (Cochran–Mantel–Haenszel test for Pan-Cancer and Fisher exact test for specific cancers, p-value <0.05, Supplementary Table 9, Fig. 5c). In two cases, i.e., *hsa-miR-320b-2* and *hsa-miR-517b*, the association of the mutations with lower cancer stages corresponded with their positive effect on patient survival. However, due to the low number of identified mutations in particular miRNA genes or hotspots, the abovementioned associations with survival and cancer stages are of very low statistical power (not corrected for multiple comparisons) and therefore must be interpreted cautiously and cannot be generalized without further experimental validation.

### KEGG pathways associated with miRNA gene mutations

Finally, to identify pathways/processes enriched in the genes regulated by the most frequently mutated miRNA genes, we used miRPath v3.0 to perform KEGG pathway enrichment analysis. As shown in Supplementary Table 10 and Fig. 6, the vast majority of the associated (adjusted p<0.01) KEGG pathways are related to different cancers or cancer-related processes, such as the cell cycle, proliferation, or apoptosis. For example, the top ten most significant associations (p<0.000005) include the following terms: Proteoglycans in cancer, Signaling pathways regulating pluripotency of stem cells, Renal cell carcinoma, Glioma, ErbB signaling pathway, Hippo signaling pathway, FoxO signaling pathway, and Wnt signaling pathway (Supplementary Table 10, Fig. 6).

**Figure 6:**
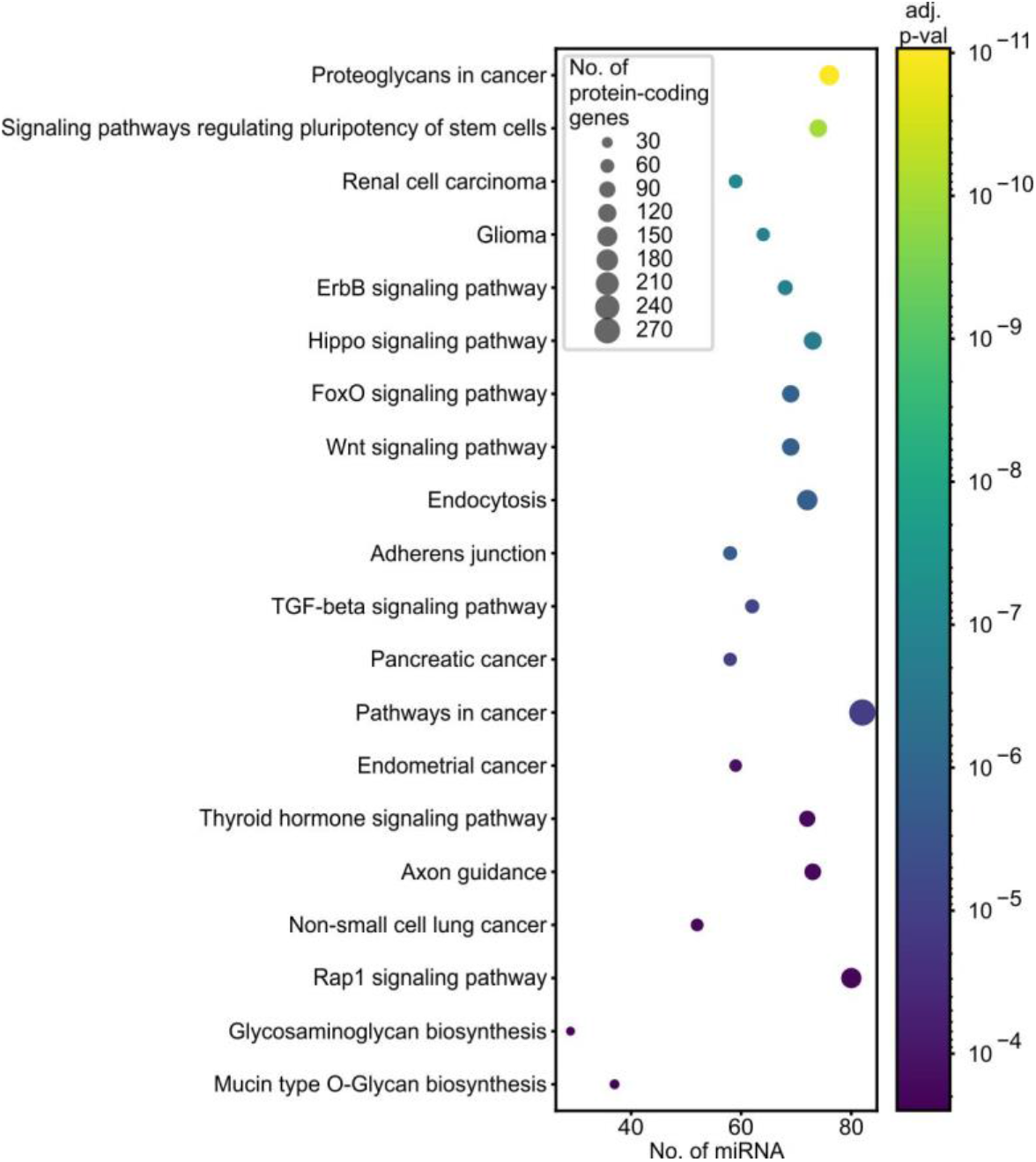
Functional association of overmutated miRNA genes with KEGG pathways. The graph shows the top 20 pathways (y-axis) enriched in protein-coding genes regulated by miRNAs (x-axis) encoded by overmutated miRNA genes. Dot size indicates the number of protein-coding genes; dot color depicts an adjusted p-value of association. The enrichment analysis was performed with the use of miRPath v3.0 and encompassed the top 100 miRNA genes enriched in the functionally weighted test. The full list of enriched pathways is shown in Supplementary Table 10.

## Discussion

Multiple functional somatic mutations with roles in cancer are known in the coding portion of the genome. In this study, we identified 7,110 mutations in miRNA genes across 33 cancer types based on data available in the TCGA repository. Most of the mutations were substitutions (~89%), with indels overrepresented within a couple of analyzed cancer types (COAD, STAD, and UCEC). Overall, approximately 33% of Pan-Cancer samples have at least one mutation in miRNA genes, with percentages substantially differing among cancer types, similar to what is observed for mutations in other genomic regions. The mutations were in general evenly distributed across miRNA gene functional subregions. This could be attributed to the fact that the majority of detected sequence variants are spontaneous mutations randomly accumulating in the cancer genome.

Among the identified mutations, we found ones located in miRNA genes playing an important and well-recognized role in cancer (e.g., *hsa-let-7* family, *hsa-miR-205* and *hsa-miR-142*) as well as miRNAs that were not yet investigated broadly in relation to cancer. In total, we identified 108 overmutated miRNA genes within the Pan-Cancer cohort and 80 overmutated miRNA genes within individual cancer types. In particular, we found multiple mutations in *hsa-miR-142, hsa-miR-205, hsa-let-7d, hsa-miR-411, hsa-miR-519e, hsa-miR-664b, hsa-miR-585, hsa-miR-496*, and *hsa-miR-1302-3*. Although the frequency of mutations in overmutated miRNA genes is lower than in commonly mutated drivers such as *TP53, CDKN2A*, or *KRAS*, it is comparable to the cancer-specific frequencies of mutations in many other protein-coding driver genes that are generally much longer than miRNA genes, e.g., *MET* (7%), *RB1* (4%), and *RIT1* (2%) in LUAD ^107^, *HRAS* (4%), *PTEN, RB1, NF1* (<1%) in PTC ^108^ or *DRD5* (3%), and *BRAF* (2%) in GBM ^109^.

Additionally, we identified 62 hotspot positions in Pan-Cancer and 69 in individual cancers, including 5 cancer-specific hotspots. One group of recurring mutations covers primarily insertions and deletions in short repeats. The mutations were identified predominantly in STAD and UCEC cancers known to be associated with MSI. It was previously suggested that simple repeats in human miRNA genes are relatively rare and preserved from mutations due to MSI ^110^. Only three such mutations were identified in hsa-miR-1303, hsa-miR-567, and hsa-miR-1273. In our study, we identified indels associated with MSI in numerous miRNA genes (e.g., *hsa-miR-320c-1*, *hsa-miR-320b-2*, and *hsa-miR-1249*), including previously observed ones. Many MSI-associated mutations are recurrently mutated hotspots; for example, *hsa-let-7d* (chr9:94178817delAA[+]) in UCEC, COAD, and Pan-Cancer (Fig. 4c), *hsa-miR-1303* (chr5:154685821TTA>T[+]) in STAD, UCEC, and Pan-Cancer and *hsa-miR-567* (chr3:112112876TA/TAAA>T[+]) in UCEC and Pan-Cancer. This result shows that the idea of the involvement of MSI in mutations within miRNA genes should be revisited, especially as those mutations may also be functional ^111,112^.

As mentioned earlier, depending on localization within the miRNA gene, mutations can have multiple effects on miRNA functionality, including changes in targets (mutations within seeds) or processing (mutations that change structure or are located in the DROSHA/DICER1 cleavage site), resulting in changed miRNA levels and/or strand balance. In our study, we detected 536 mutations and 7 recurrently mutated hotspots in seed sequences of different miRNAs, including miRNAs with defined roles in cancer. As shown before, in our previous study, such mutations affect the vast majority of predicted miRNA targets. An example of a seed hotspot mutation is chr1:209432167C>T[+] in miR-205-5p, which affects most of the predicted miRNA targets. Consistent with the putative effect of mutations on the effectiveness of miRNA biogenesis, we identified many associations of recurrently mutated genes with the level of the corresponding miRNA, predominantly resulting in decreased (e.g., miR-664b-3p, miR-134-5p) levels of the affected miRNAs. Although probably not all of the observed miRNA aberrations play any relevant functional role in cancer, a vast excess of downregulated miRNAs confirms that mutations in miRNA genes have mostly destructive effects on the structure or stability of the miRNA precursors, making them less optimal substrates for the miRNA biogenesis process. Changes in miRNA levels are a known aspect of cancer characteristics; however, they are usually attributed to other mechanisms, and the effect of somatic mutations on miRNA expression has not been systematically studied before.

Subsequent analyses of available clinical data, including patient survival and cancer stage, showed that mutations in 12 miRNA genes were associated with different metrics of patient survival (predominantly with decreases in survival), and mutations in 18 miRNA genes were associated with cancer stage. This observation further confirms the potential functionality of the miRNA gene mutations acting directly (e.g., a mutation in seed), by changes in miRNA levels, or by other secondary effects. Although we tested only the effects of overmutated miRNA genes, we cannot exclude the possibility that some of the individual mutations also affect miRNA biogenesis/function and/or cancer. On the other hand, the identified associations do not prove the functionality of the particular mutations or groups of mutations in cancer. To provide such proof, independent functional analyses are needed, in which the results presented in our study may serve as a starting point or support. Such analyses will often have to be performed in the context of a particular cancer type or condition. On the other hand, globally, the nonrandom character of the identified mutations was confirmed by a strong association of overmutated miRNA genes with KEGG pathways, of which the vast majority were specific to particular cancers or cancer-related processes.

Many approaches have been developed to discover and evaluate cancer-driver mutations in protein-coding sequences, e.g., MutSig2CV, HotSpot 3D, CLUMPS, and PARADIGM-SHIFT ^113–116^, and numerous cancer-driving mutations and genes have been identified by taking advantage of these tools. The majority of these approaches take into account (i) well-known and easy to predict consequences of mutations in protein-coding sequences, i.e., distinguishing frameshift, nonsense, missense, splicing or synonymous mutations, (ii) the predicted effects of the mutations on the AA properties and/or tolerance of AA change in particular protein domains and/or the effect of AA change on protein structure, and (iii) the conservation of the particular AA residue or particular protein. These factors allow estimation of the excess of deleterious functionally relevant mutations over neutral variants, which is one of the most important components of models identifying signals of cancer-driven selection. Unfortunately, such tools cannot be utilized for the identification of drivers in noncoding regions, including sequences encoding “noncoding” RNA. Recognizing this limitation, several approaches dedicated to the identification of drivers in noncoding sequences or with added functionalities for this purpose have been proposed (e.g., oncodriveFML, MutSigNC, ncDriver, and LARVA) ^52,55,117,118^. It was also recognized that due to different roles and functionalities, e.g., promoters, 5’ and 3’ untranslated regions (5’ and 3’ UTRs), introns, long noncoding RNAs, and miRNAs, different noncoding elements have to be analyzed with separate approaches or assumptions ^55^. Nonetheless, among the available tools, the functionality of noncoding mutations is mostly recognized by two factors: impact on protein (e.g., transcription factor) binding properties and impact on RNA structure. Sometimes, for specific ncRNA regions, additional factors are taken into account, such as the impact on miRNA binding sites in 3’UTRs. Although the structure is an important factor of miRNA biogenesis/functionality, the impact on RNA structure (e.g., the RNAsnp score) is inferred based on structures predicted for standardized tailing RNA fragments not corresponding to the size and coordinates of miRNA precursors. Therefore, at present, there is no approach/algorithm dedicated to recognize driving selection signals in miRNA genes. To overcome this limitation, in addition to evaluating the excess of the mutation in particular genes, we also weighted the mutations based on our proposed functionally related factors, with higher scores for mutations within the seed sequence, mature miRNAs, DROSHA/DICER1 cleavage sites, and disrupting protein binding motifs. Our results, together with recently published insights on mutations occurring in noncoding regions ^48,52^, may provide a basis for the development of new tools focused on miRNA cancer drivers based on described potentially functional mutations.

As there is no list of previously defined miRNA driver genes, we could not formally validate our approach; however, among the top-scored overmutated miRNA genes, we identified *hsa-miR-142*, which is the only miRNA gene in which mutations were identified in several hematologic neoplasms in several studies ^49–53,56,57^, and their cancer relevance was functionally confirmed ^27^. Our analysis confirmed the recurrence of *hsa-miR-142* mutations in hematologic neoplasms, i.e., LAML, DLBC, and also the newly identified mutation in the Burkitt lymphoma Raji cell line, but also showed mutations in several solid tumors, i.e., UCEC, BLCA, GBM, and BRCA. Additionally, thanks to the large number of mutations identified in our study and the cumulative analysis of previously detected mutations, we could illustrate for the first time the distribution of mutations in the gene. This result showed that mutations may occur in any part of the gene, not only in the seed sequence, which indicates their loss-of-function character, acting most likely by destabilizing the precursor structure and impairing miRNA biogenesis. This observation may also have the more general intriguing implication that miRNA precursors are overall quite fragile structures that may be affected by almost any mutation within the hairpin-coding sequence. Such hypotheses may be tested by the functional analysis of a higher number of randomly selected mutations in different miRNA genes.

There are several limitations of computational analyses such as the one presented in our study. First, further functional analyses of the identified recurring mutations are needed to verify their role in particular cancers. Second, not all known (miRBase) miRNA genes were covered by TCGA WES experiments. Additionally, due to different versions of WES systems used in different TCGA projects, the sequencing of some miRNA genes may not be equal in all samples. Third, even working with over 10,000 samples, the statistical power of some analyses is not sufficient, and further analyses with even larger cohorts of particular cancers or groups of cancers are awaited. Forth, some of the TCGA cancer type cohorts are quite heterogeneous, consisting of samples of different genetic background. Finally, the analyses of mutations involved in cancers would also benefit from a better understanding of the structure of miRNA genes, including more complete information about the full sequence of miRNA transcriptional units (full pri-miRNA sequences) and their regulatory elements ^119^ and better functional validation/annotation of the known miRNA genes, as proposed, e.g., in the miRGeneDB database ^38^.

In summary, we present the first comprehensive Pan-Cancer study of somatic mutations in miRNA genes in a large cohort of cancer samples. As a result, we detected thousands of different mutations located in different functionally relevant parts of miRNA genes, and many miRNA genes were overmutated either in Pan-Cancer or in specific cancer types. The frequency of the mutations in some of the overmutated miRNA genes corresponds to that observed in some validated protein-coding driver genes. Subsequent analyses (miRNA expression, survival analyses, and functional pathway associations) suggest that at least some of the overmutated miRNA genes or hotspots in miRNA genes may be driven by cancer-positive selection and therefore may play a role in cancer. Nonetheless, the functionality of particular mutations needs to be experimentally validated with the use of appropriate functional tests. Our results are also the first step (form the basis and provide the resources) for the development of computational and/or statistical approaches and tools dedicated to the identification of cancer-driver miRNA genes.

## Acknowledgments

The results published here are based upon data generated by the TCGA Research Network: https://www.cancer.gov/tcga. This work was supported by research grants from the Polish National Science Centre [2016/22/A/NZ2/00184 (to P.K.) and 2015/17/N/NZ3/03629 (to M.O.U-T.)]

## Authors contributions

MUT – participated in conceiving the study, wrote all the scripts, performed most of the computational and statistical analyses, drafted the manuscript, prepared figures, tables, and supplementary materials; PGM – participated in conceiving the study, discussed the study on all steps of analyses, participated in manuscript preparation, performed some analyses (3D structures); PN – discussed the analyses on all steps of the study, performed some analyses, participated in manuscript preparation; EK – performed the sequencing experiments, critically read and corrected manuscript; SS – performed the sequencing experiments, critically read and corrected manuscript; MG – provided samples for the *hsa-miR-1324* and *hsa-miR-142* sequencing validation experiments, advised on hematological neoplasms and head and neck cancer, critically read and corrected manuscript; PK – received financing, participated in conceiving the study, supervised and coordinated the study, drafted the manuscript (with MUT).

## Supplementary Materials

**Supplementary Table 1**. **Coordinates of 1918 miRNA genes used in the study.**

**Supplementary Table 2. All mutations identified in miRNome in the Pan-Cancer cohort with detailed characteristics.**

Each mutation within the miRNA gene is characterized by position in the genome, including strand orientation, patient TCGA ID, reference/alternative alleles detected with read-count within normal and cancer samples, mutation type and type of substitution if applicable, algorithms that detected the mutation, location of the mutation within pre-miRNA, cancer type, the balance of the miRNA strands, miRBase ‘high confidence’ label, miRGeneDB id and values used for weighted analysis (mutation within DROSHA/DICER1 cleavage region, mutation within duplex, mutation in seed, mutation disrupting protein-interacting motif). Only mutations from non-hypermutated samples are included.

**Supplementary Table 3. Lists and characteristics of overmutated miRNA genes identified in Pan-Cancer and particular cancer types by (A) ordinary binomial and (B) functionally weighted mutation distribution analysis.**

**Supplementary Table 4. Percentage of mutations in overmutated miRNA genes in individual cancer types and Pan-Cancer.**

The table shows the exact values of the frequencies visualized in Figure 3B.

**Supplementary Table 5. List and characteristics of significant hotspots in miRNA genes identified in Pan-Cancer and in particular cancer types.**

**Supplementary Table 6. List of mutations detected in *hsa-miR-142* in previous studies.** Hg38 genome positions were assigned regardless of what genome version was used in a study.

**Supplementary Table 7. List and characteristics of miRNAs whose expression levels are associated with mutations in the corresponding miRNA genes.**

MW - Mann-Whitney test.

**Supplementary Table 8. List and characteristics of overmutated miRNA genes associated with patient survival (PFI, DFI, DSS, and OS).**

**Supplementary Table 9. List and characteristics of overmutated miRNA genes associated with cancer stage.**

**Supplementary Table 10. Significantly enriched KEGG pathways for the top mutated miRNA genes in Pan-Cancer analyzed with miRPath v3.0.**

The analysis was performed for 100 mature miRNAs encoded by the most significantly overmutated miRNA genes based on the functionally weighted analysis.

**Supplementary Figure 1:**
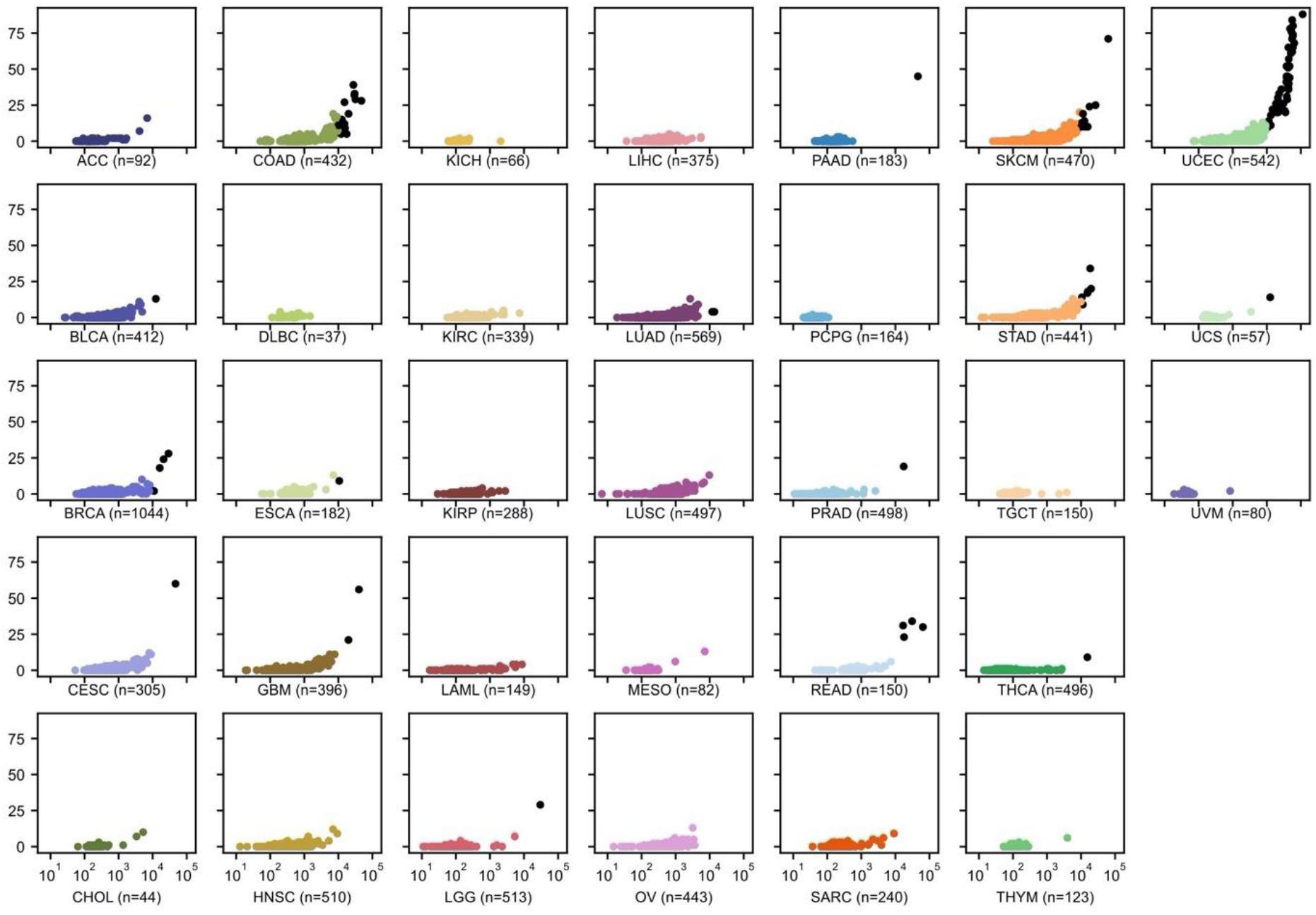
The number of mutations in the exome (x-axis, log_10_ scale) and miRNome (y-axis, linear scale) per sample in each of the analyzed cancer types. The X-axis was unified for all plots. Each dot represents a single sample. Black dots represent hypermutated (>10k somatic mutations in exome) samples.

**Supplementary Figure 2:**
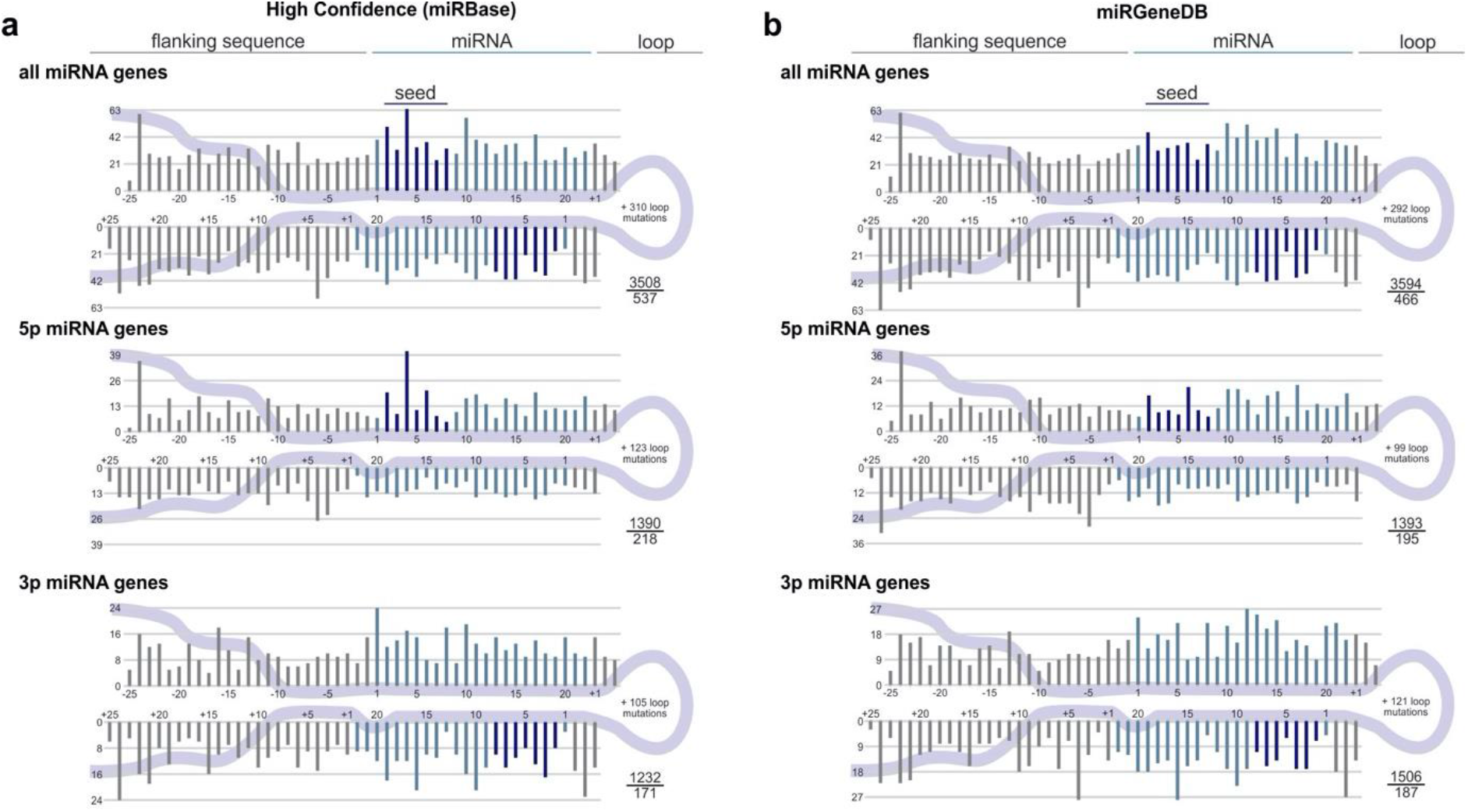
Localization of somatic mutations in miRNA genes annotated as (a) ‘high confidence’ in miRBase and (b) mirGeneDB. The scheme of the figure is as shown in Fig. 2.

**Supplementary Figure 3:**
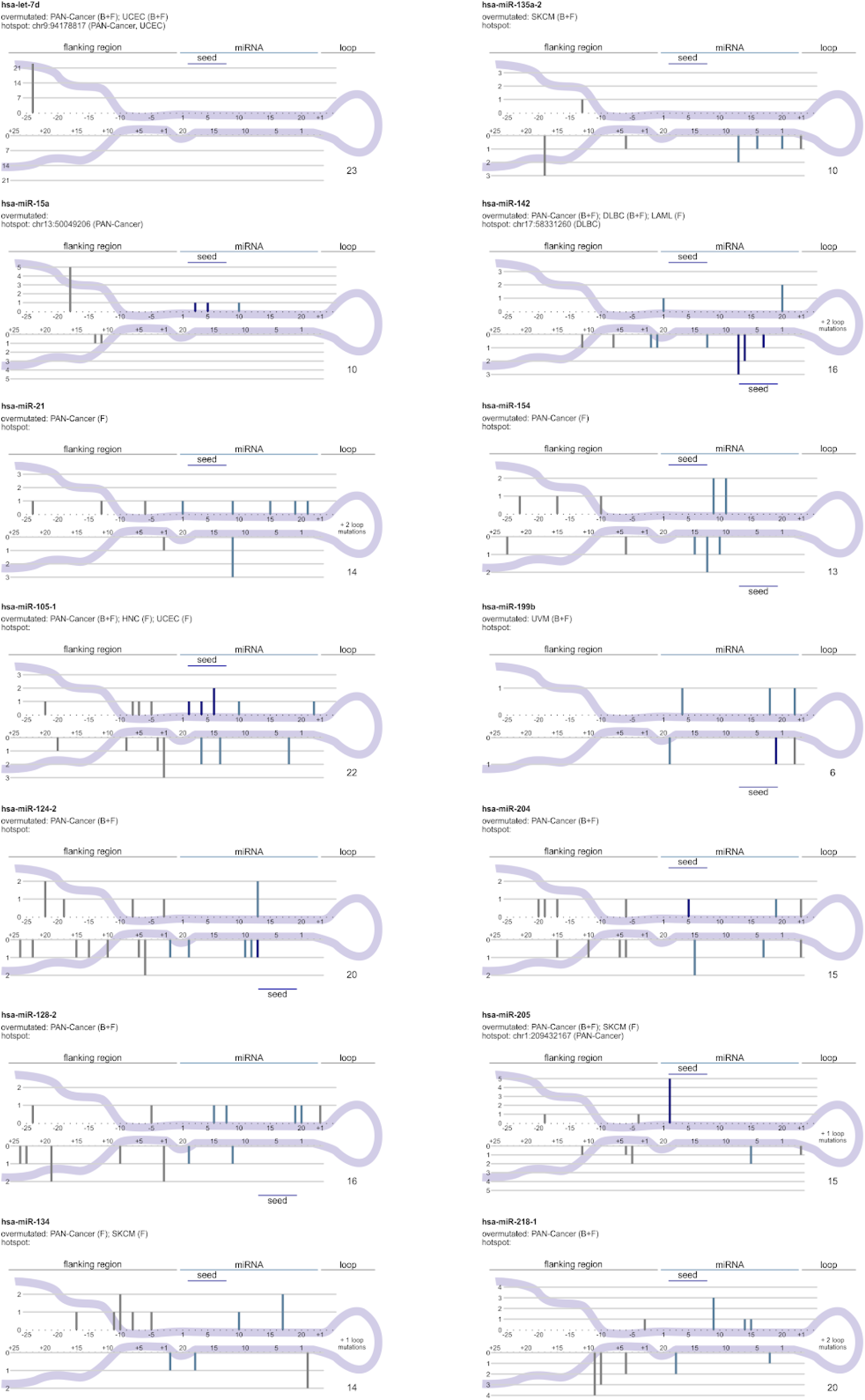
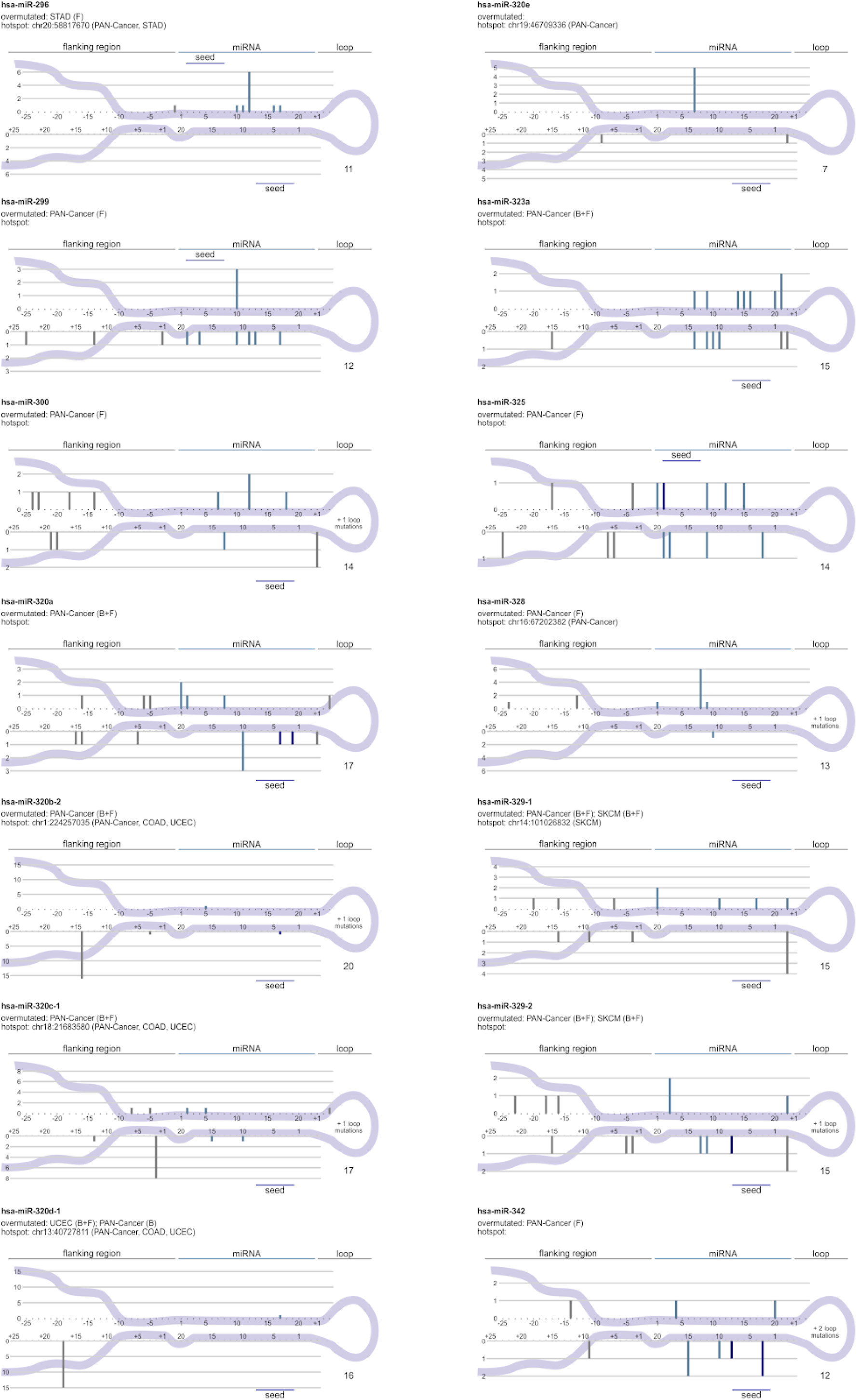
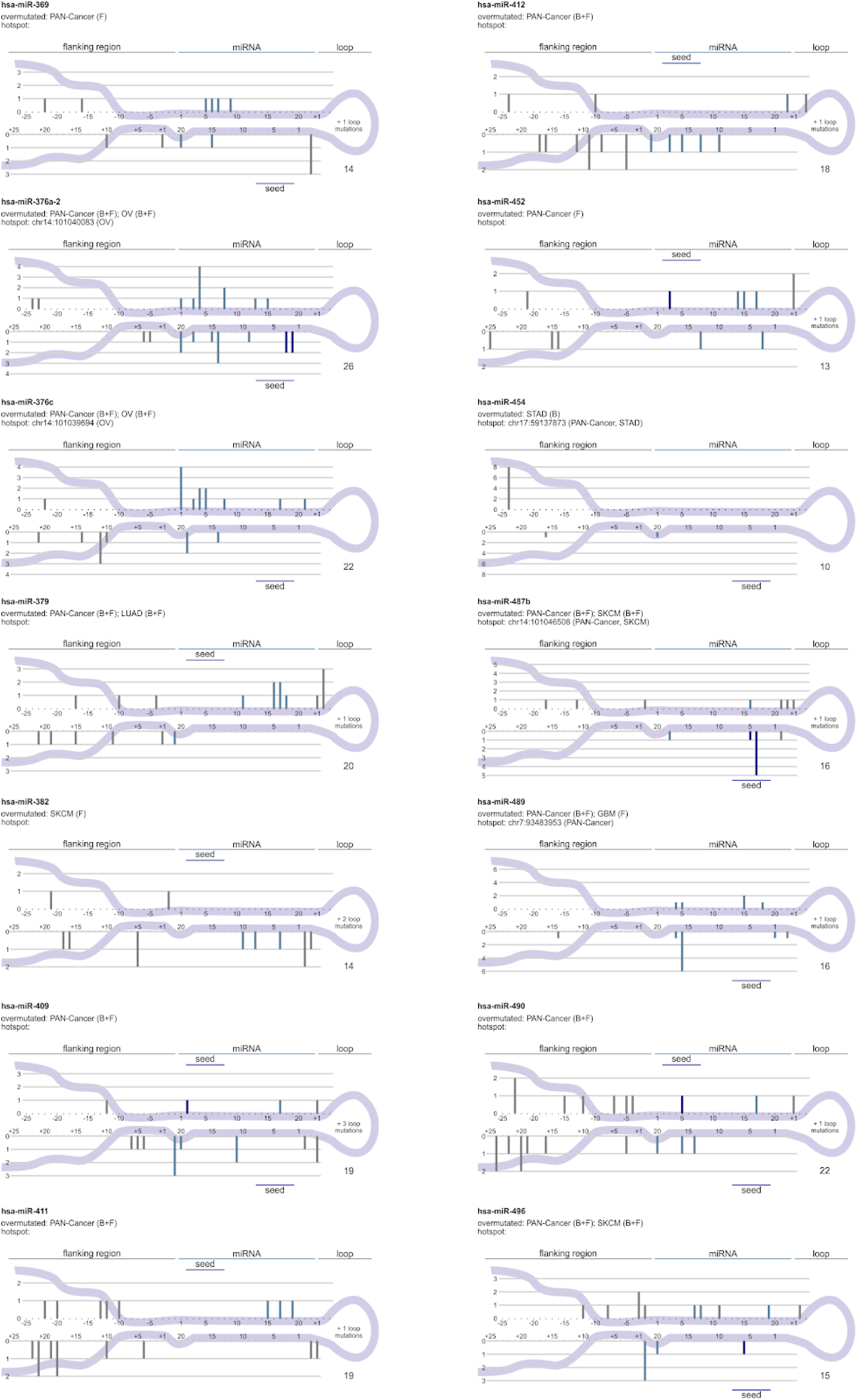
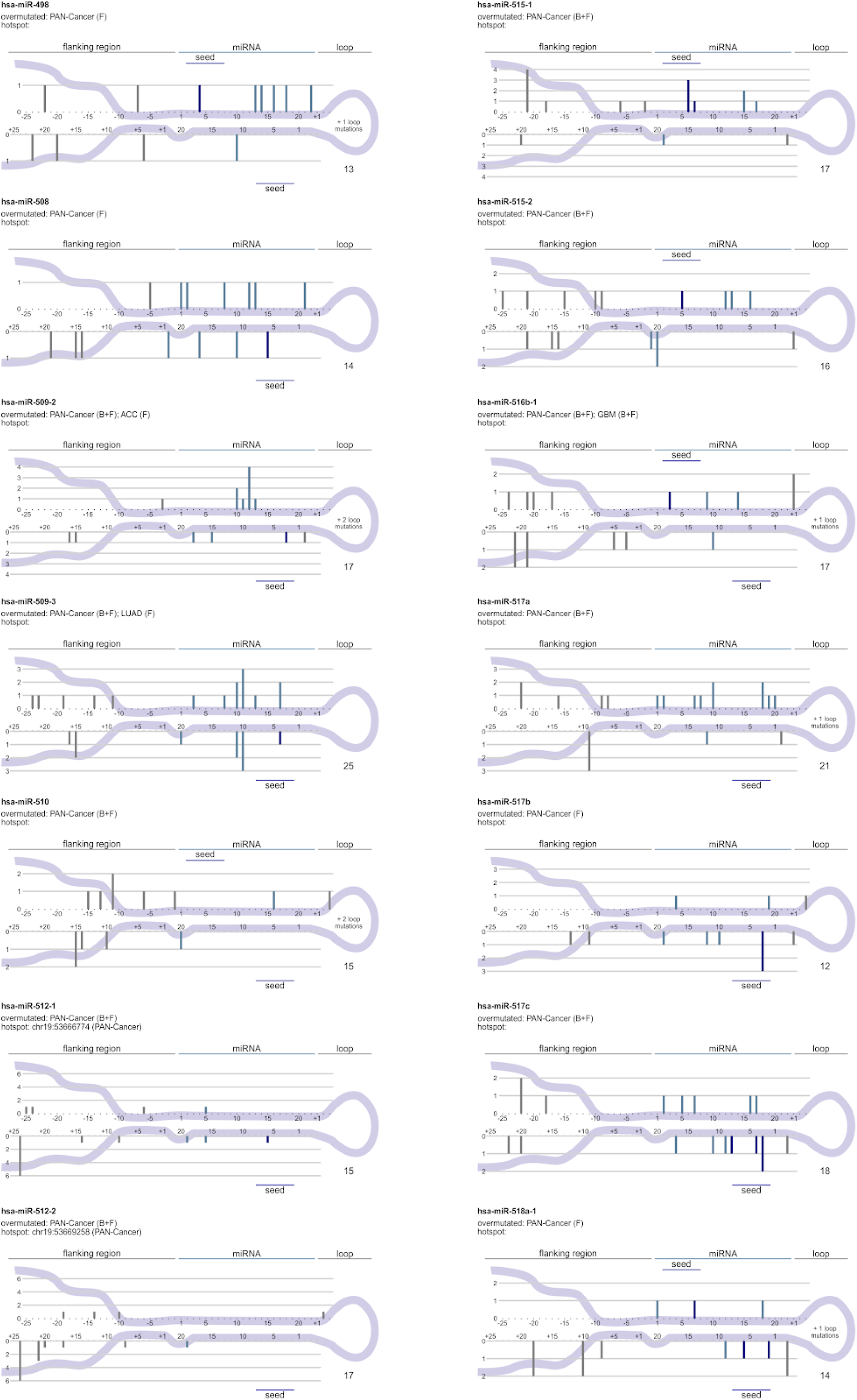
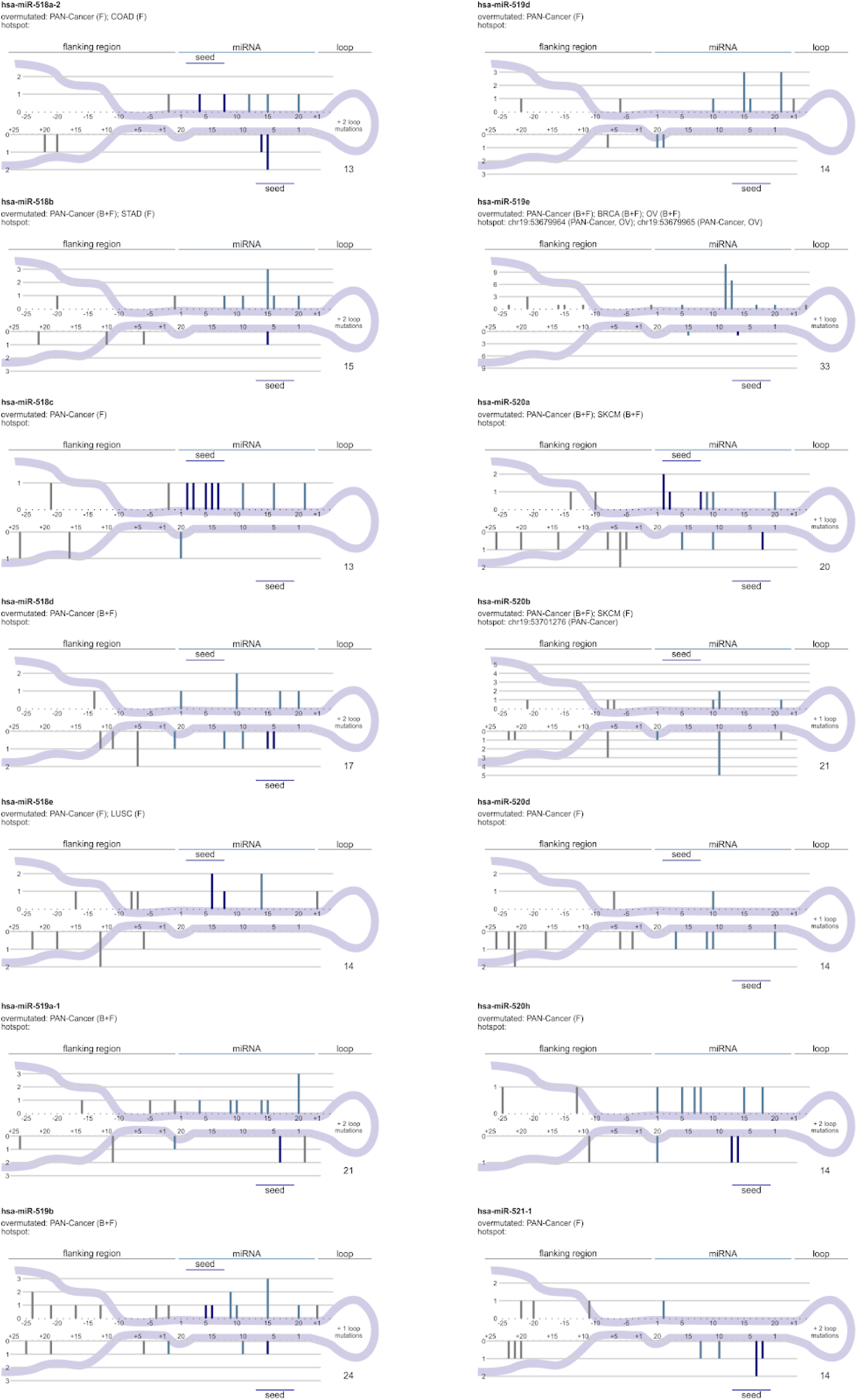
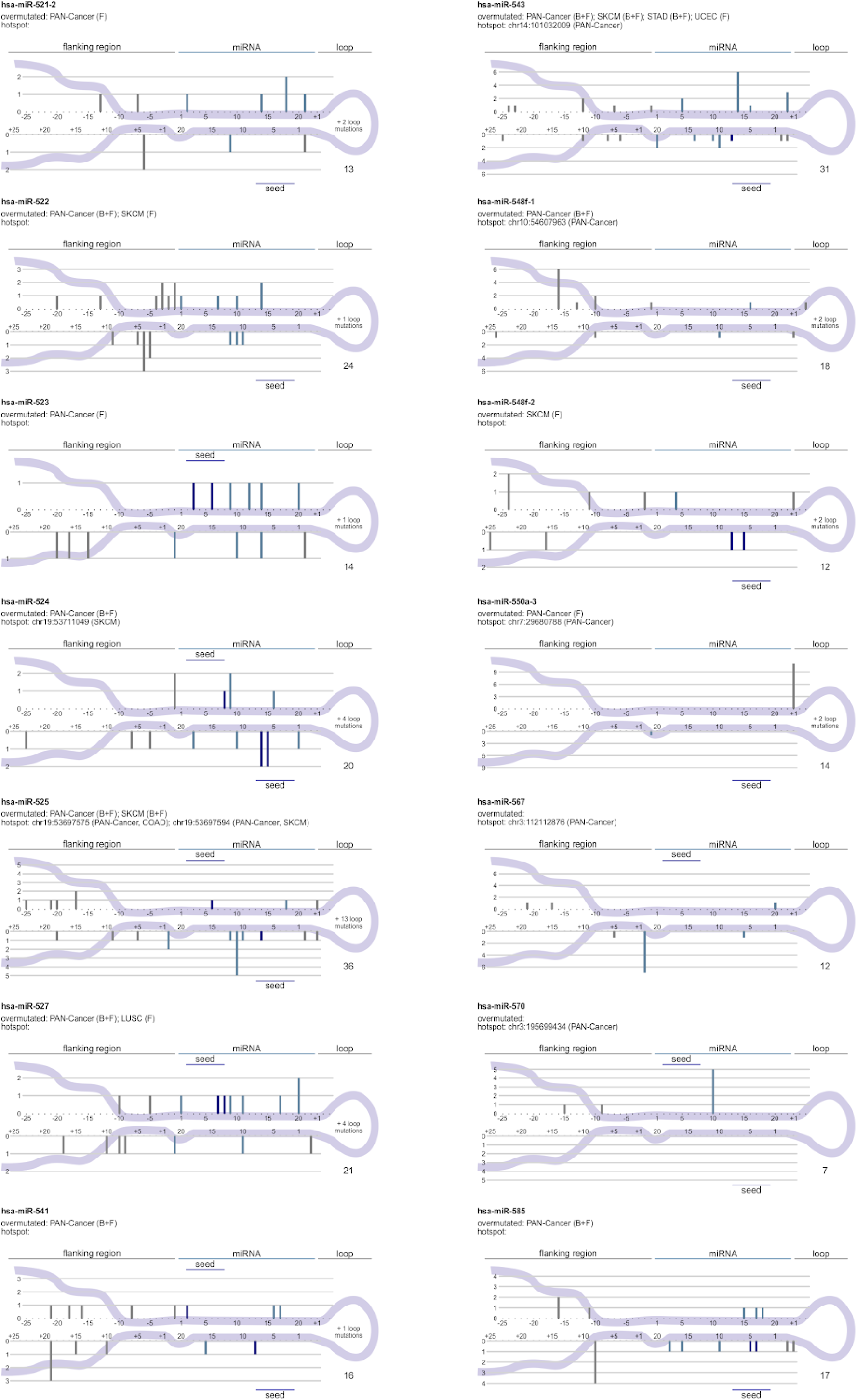
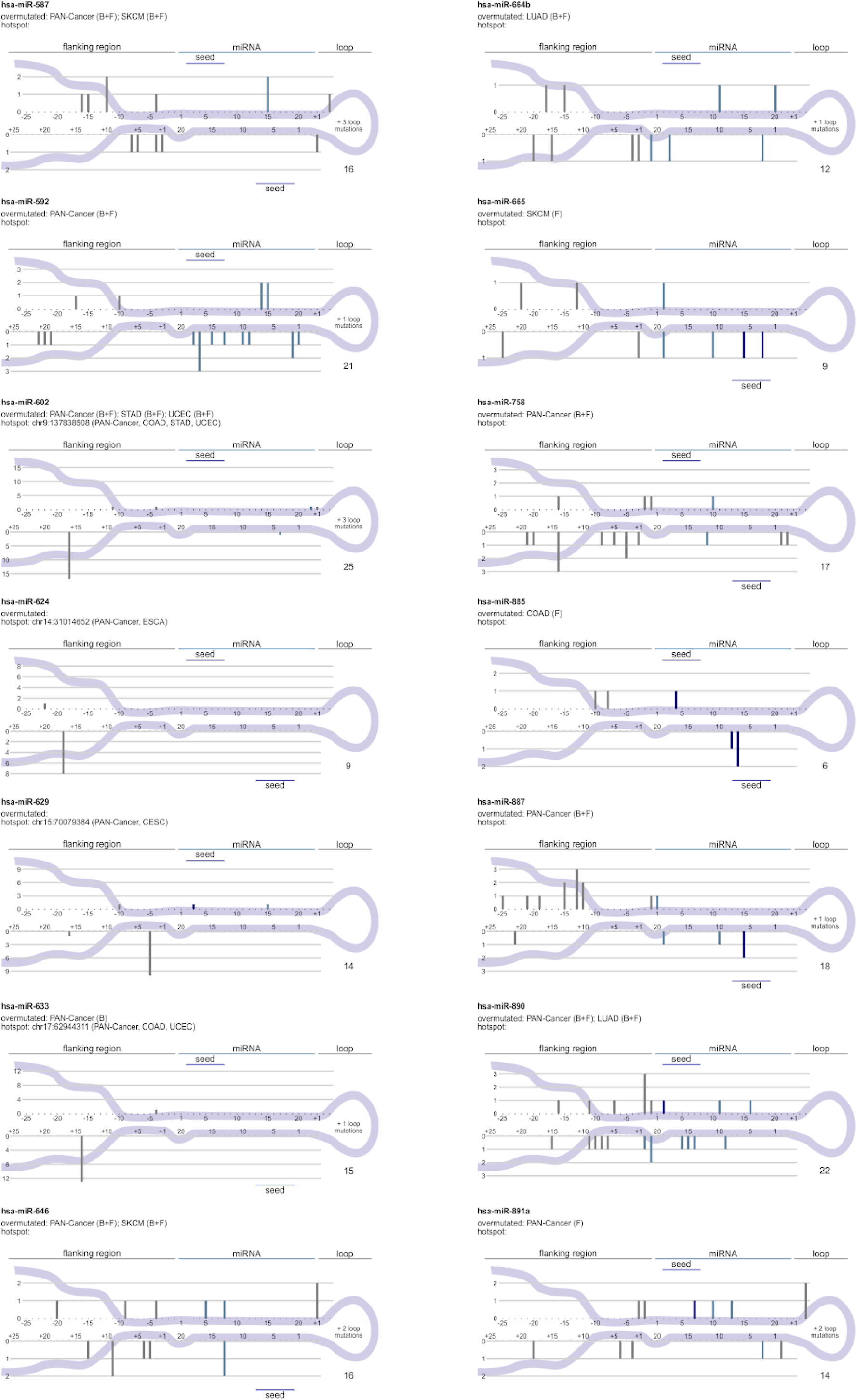
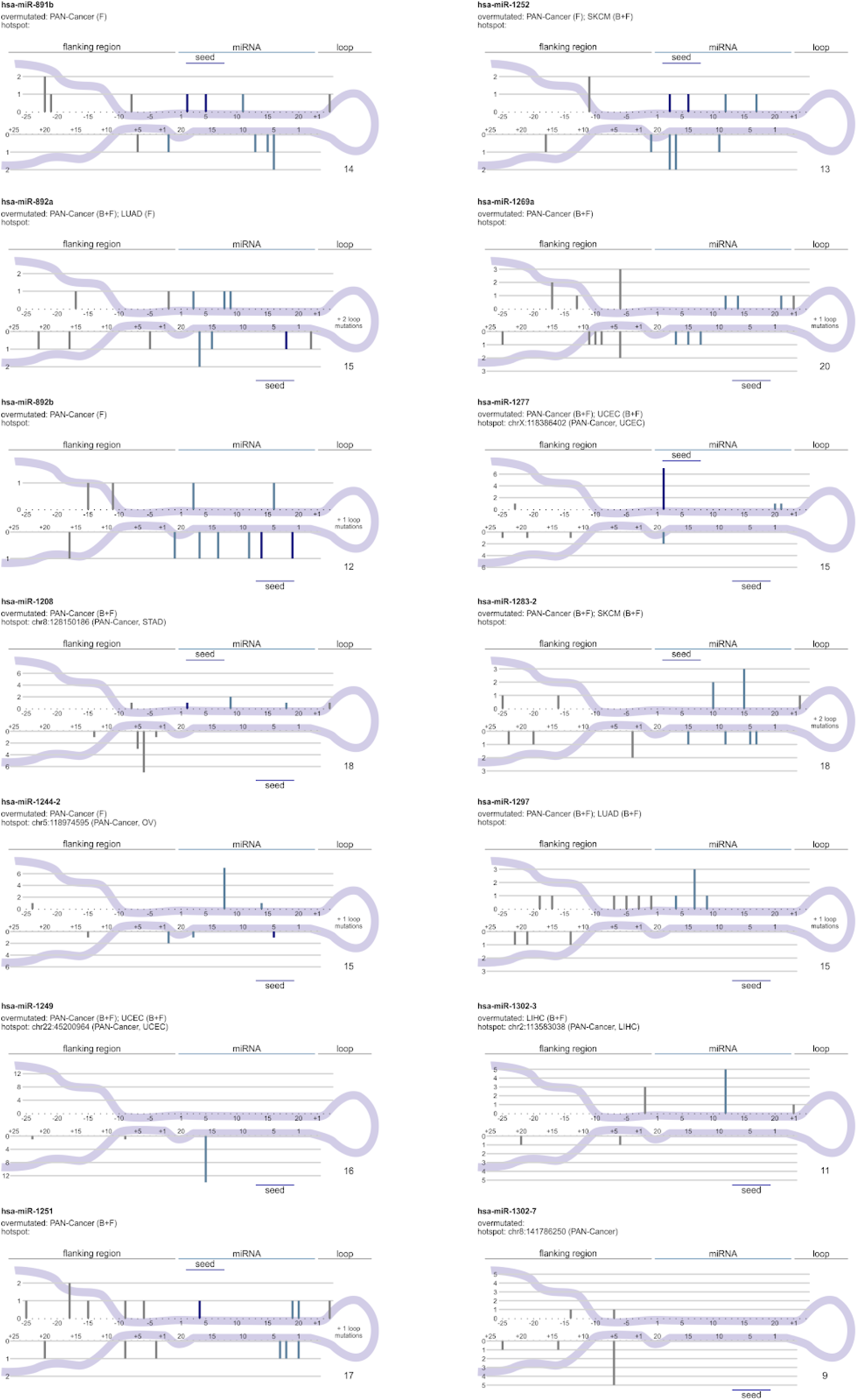
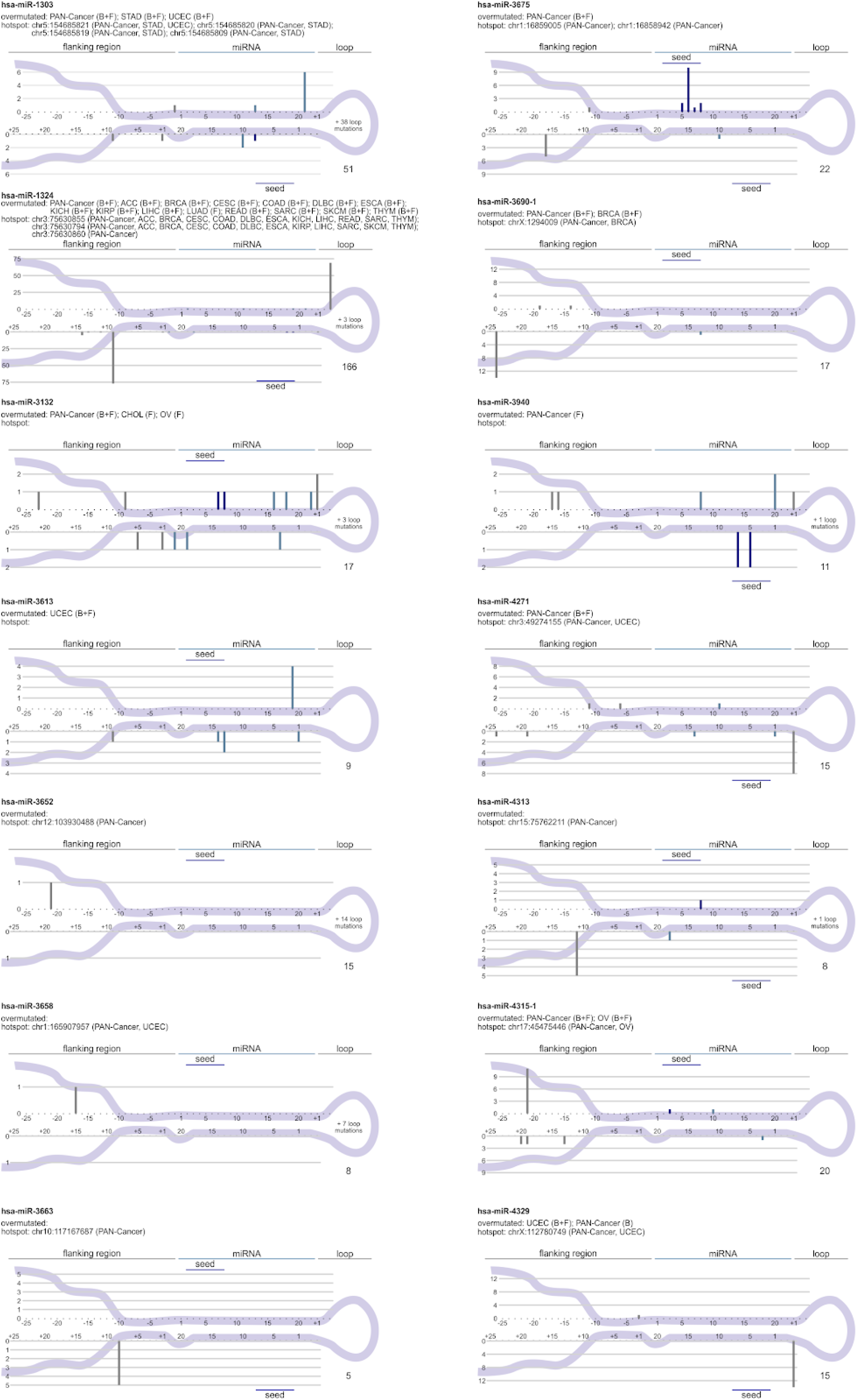
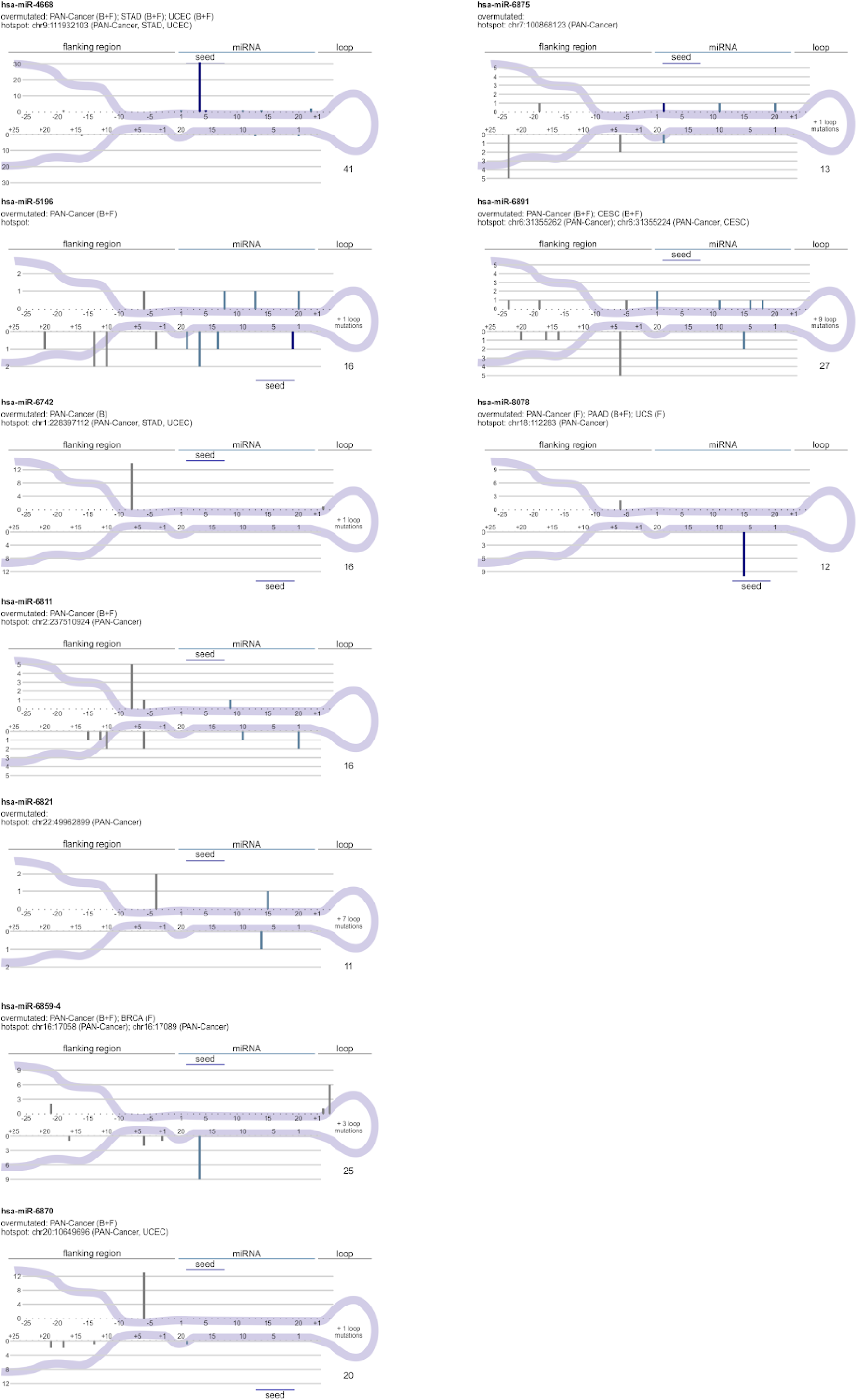
Localization of mutations in the miRNA genes (136) either overmutated or containing hotspot mutations. B and F indicate ordinary binomial and functionally weighted analyses, respectively. The numbers in the right lower corner indicate the total number of mutations.

